# Therapeutic efficacy of an immune stimulatory gene therapy strategy in a mouse model of high grade brainstem glioma

**DOI:** 10.1101/846097

**Authors:** Flor M. Mendez, Padma Kadiyala, Felipe J. Nunez, Stephen Carney, Fernando Nunez, Jessica C. Gauss, Ramya Ravindran, Sheeba Pawar, Marta Edwards, Pedro R. Lowenstein, Maria G. Castro

## Abstract

**Purpose:** Diffuse intrinsic pontine glioma (DIPG) bears a dismal prognosis. A genetically engineered brainstem glioma model harboring the recurrent DIPG mutation, ACVR1-G328V (mACVR1), was developed for testing an immune-stimulatory gene therapy.

**Experimental Design:** We utilized the Sleeping Beauty transposase system to generate an endogenous mouse model of mACVR1 brainstem glioma. Histology was used to characterize and validate the model. We performed RNAseq analysis on neurospheres (NS) harboring mACVR1. mACVR1 NS were implanted into the pons of immune competent mice to test the therapeutic efficacy and toxicity of immune stimulatory gene therapy using adenoviruses expressing thymidine kinase (TK) and fms-like tyrosine kinase 3 ligand (Flt3L). mACVR1 NS expressing the surrogate tumor antigen ovalbumin were generated to investigate if TK/Flt3L treatment induces the recruitment of tumor-antigen specific T cells.

**Results:** Histological analysis of mACVR1 tumors indicates that they are localized in the brainstem and have increased downstream signaling of bone morphogenetic pathway as demonstrated by increased phospho-smad1/5 and Id1 levels. Transcriptome analysis of mACVR1 NS identified an increase in the transforming growth factor beta (TGF-β) signaling pathway and the regulation of cell differentiation. Adenoviral delivery of TK/Flt3L in mice bearing brainstem gliomas resulted in anti-tumor immunity, recruitment of anti-tumor specific T cells and increased median survival.

**Conclusions:** This study provides insights into the phenotype and function of the tumor immune microenvironment in a mouse model of brainstem glioma harboring mACVR1. Immune stimulatory gene therapy targeting the hosts’ anti-tumor immune response inhibits tumor progression and increases median survival of mice bearing mACVR1 tumors.

**Translational Relevance:** The therapeutic efficacy of anti-DIPG immune responses is limited due to a low number of immune cells in the tumor microenvironment. We have uncovered a novel treatment strategy that utilizes adenoviral delivery of therapeutic genes, thymidine kinase (TK) and fms tyrosine kinase 3 ligand (Flt3L) into the tumor, eliciting a reprograming of the host’s own immune system to recognize and kill tumor cells. We demonstrate that TK/Flt3L therapy generates an effective anti-tumor response and can be safely delivered into the brainstem. This treatment approach could provide a novel translational approach towards potentiating an anti-DIPG immune response to overcome the current limitations in the treatment of patients with DIPG.

## Introduction

Diffuse intrinsic pontine glioma (DIPG) is an aggressive, diffusive brain tumor that originates in the pons and is diagnosed based on clinical and radiological criteria (1). DIPG occurs mostly in children of median age 6-7 years; it is an inoperable brain tumor and despite multiple trials testing various chemotherapeutic strategies, none have demonstrated a survival benefit. Thus, focal radiation remains the standard of care (1,2). The median overall survival for DIPG is 10.8 months, and the 2 year survival is ~5.2% (3). In 2014, four independent studies that comprised data from 195 DIPGs, identified Activin A receptor type I (ACVR1) as the most recurrently mutated gene following the histone H3 K27M mutation (4-7). Six recurrent, somatic mutations in ACVR1 were found in 24% of DIPG cases (8). The characteristics of DIPG patients presenting with the ACVR1 mutation include younger age of onset (~5 years of age) and longer overall survival time (4,9). Sequencing revealed that DIPG tumors have a lower mutation frequency compared to adult glioblastoma, and that classical tumor pathways such as the PI3K and p53 pathways are also altered in these tumors (10).

ACVR1 encodes a type 1 serine/threonine kinase that is part of the transforming growth factor beta (TGF-ß) family. It is noteworthy to mention that ACVR1 mutations are only found in DIPG tumors; ACVR1 is not mutated in any other cancers. Thus, uncovering the role of this gene mutation in DIPG tumorigenesis and disease progression could provide novel mechanistic insights useful to develop novel therapeutic targets.

Since DIPGs are not resectable and they are highly invasive, it would be advantageous to harness the power of the immune system to elicit effective anti-tumor immunity in DIPG patients. Immune mediated treatment modalities have yielded promising clinical benefits in melanoma, non-small-cell lung cancer, renal cell cancer, and prostate cancer (11-14). We have also previously shown the efficacy of an immune stimulatory gene therapy approach in several rat and mouse models of adult glioblastoma (15,16). This approach has recently completed its Phase I Clinical Trial accrual for the treatment of adult patients with newly diagnosed glioblastoma multiforme (World Health Organization (WHO) grade IV (NCT01811992). This immune-mediated gene therapy approach is based on adenoviral delivery of herpes simplex virus type 1-thymidine kinase (TK) and Fms-like tyrosine kinase 3 ligand (Flt3L). Upon administration of the prodrug, ganciclovir, proliferating tumor cells expressing TK undergo immunogenic cell death(15,16). Dying tumor cells, release damage-associated molecular patterns (DAMPs), such as high-mobility group B1 protein (HMGB1), calreticulin, and adenosine triphosphate (ATP) (15,17). Meanwhile, Flt3L elicits the recruitment of dendritic cells to the tumor microenvironment(15). HMGB1 released by dying tumor cells activates dendritic cells through Toll-like receptor 2 (TLR2) mediated signaling(15). Activated dendritic cells pick up the tumor antigens and traffic to the draining lymph nodes, where they generate a specific anti-tumor cytotoxic T-cell response (15,18). Herein, we aimed to test the efficacy of this immune stimulatory approach in an immunocompetent mouse model of mutant ACVR1 brainstem glioma.

To test the efficacy of this immune-mediated gene therapy approach in syngeneic mouse brainstem glioma models, we developed a genetically engineered mouse model encoding mutant ACVR1 using the Sleeping Beauty (SB) transposase system(19-22). SB is a transposase that is able to recognize inverted repeats/direct repeats (IR/DR) sites on DNA transposons and carry out a cut and paste reaction to integrate transposon DNA into a host genome(23). The SB system can be utilized to generate endogenous tumors that resemble gliomas through delivery of DNA transposons that encode oncogenes and tumor suppressors (19-22,24). Transcriptome analysis of ACVR1 mutant neurospheres identified an increase in the TGF-β signaling pathway and signaling pathways regulating the pluripotency of stem cells, and a decrease in the focal adhesion pathway. Treatment with TK/Flt3L immune stimulatory gene therapy significantly improved median survival compared to standard of care. The TK/Flt3L gene therapy induced a strong anti-tumor cytotoxic immune response demonstrated by an increase in the frequency of tumor antigen-specific CD8 T cells in mice treated with TK/Flt3L therapy when compared to saline controls. Our results suggest that immune-mediated gene therapy could be a promising therapeutic approach for DIPG.

## Materials and Methods

### Experimental model

All animal studies were conducted according to guidelines approved by the Institutional Animal Care and Use Committee at the University of Michigan. Animals were housed in an AAALAC accredited animal facility and had constant access to food and water; they were monitored daily for tumor burden. Males and females were used. The strain of mice utilized in the study was C57BL/6 (Jackson Laboratory, 000664).

A murine model of brainstem glioma was generated by employing the Sleeping Beauty (SB) transposon system to integrate plasmid DNA into the genome of postnatal day 1 (P1) mouse pups. The plasmids utilized were as follows: (i) SB Transposase and luciferase (pT2C-LucPGK-SB100X, henceforth referred to as SB/Luc) (ii) a short-hairpin against p53 (pT2-shp53-GFP4, henceforth referred to as shp53) or shp53-NO-GFP (iii) a constitutively active mutant of NRAS (pT2CAG-NRASV12, henceforth referred to as NRAS) with or without (4) mutant ACVR1 G328V (pkt-ACVR1-G328V-IRES-Katushka; henceforth referred to as mACVR1)(19,20,22). To create the mACVR1 plasmid we cloned pCMV5-ALK2-WT into pKT2-IRES-Katushka by blunt cloning. Then, we used the QuikChange II Site-Directed Mutagenesis Kit (Agilent, 200523) to introduce (c.983G>T, p.Gly328Val) mutation into pKT2-ALK2-WT-IRES-Katushka, to generate pKT2-ACVR1-G328V-IRES-Katushka (Addgene plasmid #77437). The resultant mACVR1 plasmid was confirmed by Sanger sequencing. SB/Luc, shp53 and NRAS plasmids were the generous gift of Dr. John Ohlfest (University of Minnesota, now deceased). The pCMV5-ALK2-WT plasmid was a generous gift from Jeff Wrana (Addgene plasmid #11741). All experiments were performed using post-natal day 1 (P1) or P2 wild-type C57BL/6 mice. The plasmid combinations injected were as follows: (1) (i) shp53 and NRAS (henceforth referred to as wt-ACVR1), (ii) shp53, NRAS, and ACVR1m. Mice were injected according to a previously described protocol (20) and described in detail in the Supplementary Methods.

### Immunohistochemistry (IHC) of paraffin embedded brains

Immunhistochemistry staining was performed as previously described (20,22) and detailed in the Supplementary Methods. Antibody information is available in Supplementary Table S1.

### Primary Neurospheres (NS)

Mouse neurospheres (NS) were generated from tumors that were developed using the SB system by injection of the following plasmid combinations (1) (i) shp53 and NRAS, or (ii) shp53, NRAS, and ACVR1m into the lateral ventricle (1.5 mm AP, 0.7 mm lateral, and 1.5 mm deep from the λ-suture) following previously described protocols and detailed in the Supplementary Methods (20-22,25). Western blot analysis and inhibitor treatment with LDN-214117 is also detailed in the Supplementary Methods and antibody information is available in Supplementary Table S1. SU-DIPG-VI and SU-DIPG-XX1 were obtained from Dr. Michelle Monje at Stanford University (Stanford, Ca) in accordance with an institutionally approved protocol at each institution. Experimental details of immunocytochemistry on DIPG cultures is available in the Supplementary Methods.

### RNA-seq

RNA-sequencing was performed in collaboration with the University of Michigan sequencing core. Total RNA was isolated from tumor NS using the RNeasy Plus Mini Kit (Qiagen, 74134) and 100 ng of purified RNA were sent for analysis. The libraries were prepared using RiboGone (Takara, 634836) and TruSeq Stranded Total RNA Human/Mouse/Rat (Illumina, 20020596) with 100 ng of input and 13 PCR cycles. Sequencing was performed by the UM DNA Sequencing Core, using the Illumina Hi-Seq platform. RNA-seq analysis is available in the Supplementary Methods.

### Implantable syngeneic murine brainstem glioma models

Female C57BL/6 mice between 6-8 weeks were used for all implantation experiments. Intracranial tumors were generated by stereotaxic injection of 1000 ACVR1m tumor NS into the pons using a 5 µl Hamilton syringe with a removable 33-gauge needle with the following coordinates: (0.8mm posterior; 1.00mm lateral to the λ-suture and 5mm deep). Animals were anesthetized, then the skin over the incision site was cut and retracted, and a burr hole was drilled into one side of the skull using a 0.45 mm drill bit corresponding to the pons coordinates. Tumor NS were delivered in a 2 µl volume after holding the needle in place for 2 minutes. Each injection was performed over the course of 7 minutes; the needle was left in place for an additional minute before being slowly withdrawn from the brain. Intra-tumoral injection of adenoviral vectors and radiation treatment, flow cytometry, T cell proliferation analysis, and the cytotoxic T cell assay are detailed in the Supplementary Methods. Details of complete blood cell count and serum chemistry analysis, neuropathological analysis, and H&E staining of liver sections are elaborated in the Supplementary Methods.

## Statistical Analysis

All data were analyzed using GraphPad Prism version 8, or R (version 3.1.3). All animal studies were carried out with at least 3 animals per group (specified in each experiment). The statistical test used is indicated in each figure. A p ≤ 0.05 was considered significant.

## Results

### Brainstem gliomas induced by transforming neural progenitor cells using the Sleeping Beauty (SB) transposase system

To assess the impact of activating ACVR1 mutations in brainstem glioma pathogenesis and response to therapeutics, we generated a genetically engineered mouse model of brainstem glioma using the Sleeping Beauty (SB) transposase system (19,20). We induced brainstem tumors by activation of the receptor tyrosine kinase (RTK)-RAS-phosphatidylinositol 3-kinase (PI3K) pathway, which is upregulated in a large percentage of DIPGs, and through inactivation of *TP53*, also commonly mutated in DIPG (3,26,27). This was achieved through the delivery of the following plasmids: *NRASV12*, a short hair pin targeting tumor *protein p53* (*TP53)* (shP53), and SB transposase/firefly luciferase, with or without *ACVR1*^G328V^ (Fig.1A, Supplementary Fig. 1S) into the fourth ventricle of neonatal mice. We used bioluminescence imaging to monitor transfection efficiency and tumor development (Fig.1B). The two experimental groups were: (1) Wild-type ACVR1 (wt-ACVR1) [*NRASV12* and shp53) and (2) mutant ACVR1 (mACVR1) (*NRASV12*, shp53, *ACVR1*^G328V^). The median survival (MS) of mice in the mACVR1 group was 127 days post injection (dpi) (Fig. 1C). The median survival for the wt-ACVR1 group was MS=85 dpi (Supplementary Fig. 2S). All tumors, regardless of ACVR1 mutation status, displayed high cellularity, nuclear atypia, invasive features, and grew in the brainstem (Fig. 1D). All tumors expressed oligodendrocyte transcription factor 2 (Olig2) (Fig. 1E), glial fibrillary acidic protein (GFAP) (Fig. 1F), nestin (Fig. 1 G) and transcription factor Sox2 (Fig. 1 H). Both wt-ACVR1 and mACVR1 tumors were positive for the proliferative marker, Ki67 (Fig. 1 I), phosphorylated extracellular signal-regulated kinase (pERK) 1/2 (Fig. 1J) and phospho-MEK1/2 (Supplementary Fig. S3). Expression of pERK 1/2 (Supplementary Fig. S4A) and pMEK1/2 (Supplementary Fig. S4B) was also confirmed on human DIPG cultures: SU-DIPG VI (wt-ACVR1) and SU-DIPG XX1 (mACVR1).

**Figure 1.**
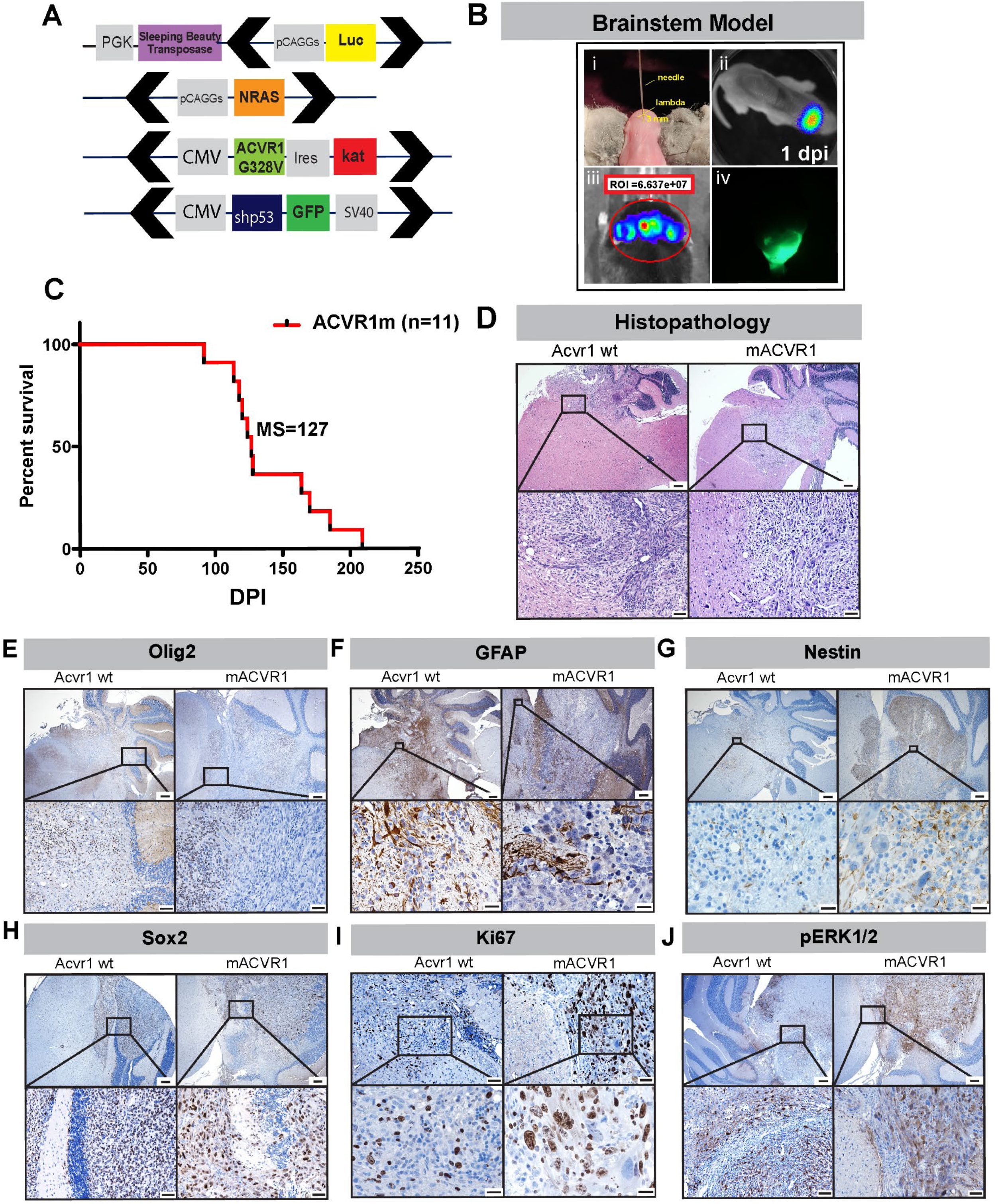
Generation of mutant (mAcvr1) mouse brainstem glioma model using the Sleeping Beauty (SB) transposase system. **(A)** Schematic representation of SB transposase and transposon plasmids used to develop gliomas in the brainstem. Black arrows indicate position of inverted repeat and direct repeat (IR/DR) sequences which flank specific DNA sequences (Luciferin, NRAS GV12, shP53-GFP, ACVR1 G328V-IRES-Katushka). **(B)** Schematic representation of brainstem glioma model. (i) One day old pups were injected in the fourth ventricle (3 mm posterior to the λ-suture and 3 mm deep) with plasmid cocktail. (ii) Bioluminescence imaging at 1-day post injection (dpi) confirming the efficiency of the in vivo transfection. (iii) Bioluminescence imaging obtained when mice displayed signs of neurological deficits confirms the presence of a large tumor. (iv) Fluorescence image of GFP^+^ brainstem tumor. **(C)** Kaplan-Meier survival curve for genetically engineered mice bearing mACVR1 (n= 11) tumor; MS: median survival. **(D)** Hematoxylin and eosin stained paraffin embedded SB brainstem tumor sections (wt-ACVR1 and mACVR1). Scale bar in the upper panel images is 200 µm. Scale bar in the bottom panel images is 20 µm. **(E-J)** Immunohistochemistry (IHC) staining for: **(E)** Olig2, an oligodendrocyte marker (scale bar in the upper panel image is 200 µm, scale bar for the bottom panel images is 50 µm) **(F)** GFAP, an astrocyte marker (scale bar in the upper panel image is 200 µm, scale bar for the bottom panel images is 20 µm) **(G)** Nestin, a neural stem cell marker (scale bar in the upper panel image is 200 µm, scale bar for the bottom panel images is 20 µm) **(H)** Sox2, a transcription factor that plays an important role in the maintenance of neural stem cells (scale bar in the upper panel image is 200 µm, scale bar for the bottom panel images is 50 µm) **(I)** Ki67, a marker of proliferating cells (scale bar in the upper panel image is 200 µm, scale bar for the bottom panel images is 20 µm) and **(J)** phosphorylated ERK-1/2 protein (pERK1/2) (scale bar in the upper panel image is 200 µm, scale bar for the bottom panel images is 50 µm).

**Figure 2.**
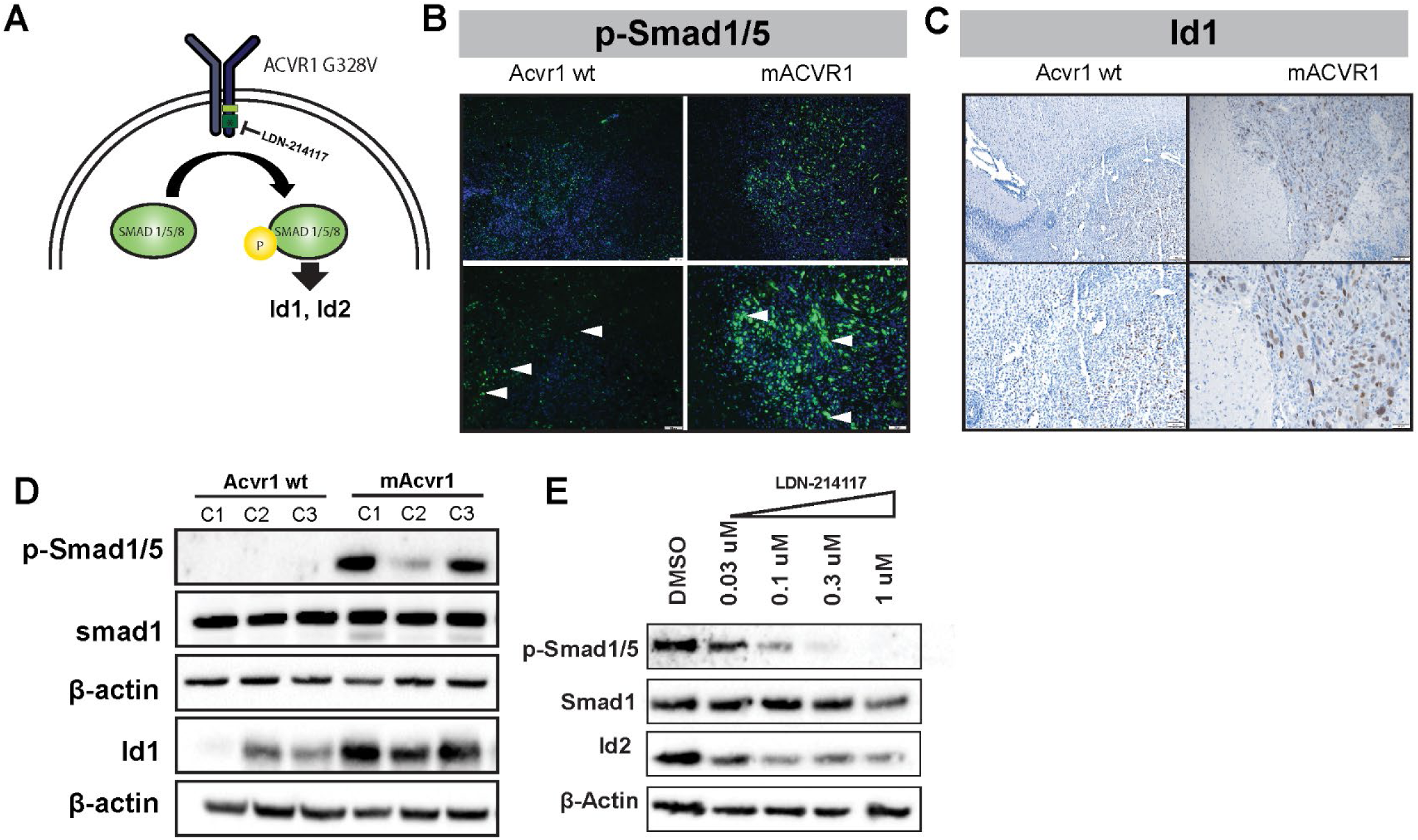
ACVR1^G328V^ activates the TGF-ß /Smad signaling pathway *in vivo* and *in vitro*. **(A)** Diagram representing the TGF-ß/Smad signaling pathway. When ACVR1 is mutated this pathway is constitutively active regardless of the presence or absence of BMP ligand. LDN-214117 inhibits ACVR1 kinase activity. **(B)** Immunofluorescence (IF) staining for phospho-Smad1/5 in paraffin embedded SB brainstem tumor sections (wt-ACVR1 and mACVR1). Scale bar in the upper panel image is 100 µm, scale bar for the bottom panel images is 50 µm. White arrows indicate positive expression. **(C)** Immunohistochemistry (IHC) staining for Id1 protein in paraffin embedded SB brainstem tumor sections (wt-ACVR1 and mACVR1). Scale bar in the upper panel image is 100 µm, scale bar for the bottom panel images is 50 µm. **(D)** WB assay was performed on three different clones of wt-ACVR1 and mACVR1 NS to assess the protein level of phopho-Smad1/5, Smad1, and Id1, a downstream Smad1/5 regulated gene. ß-actin: loading control. C1, C2, C3 represent the clones of NS used. **(E)** Western blot assay was performed on mACVR1 NS treated with LDN-214117 **(0.03-1uM)** and blotted for Smad1/5, Smad1, and Id2; ß-actin: loading control.

### mACVR1 brain stem gliomas exhibit elevated levels of phosphorylated Smad1/5

We next investigated whether tumors encoding mutant ACVR1 exhibited activated Smad1/5/8 transcription factors, downstream mediators of ACVR1 signaling (Fig. 2A). We observed that, indeed, mACVR1 brainstem gliomas displayed elevated levels of phosphorylated (phospho)-smad1/5 (Fig. 2B). This correlated with increased levels of the downstream canonical target gene, inhibitor of DNA binding 1 (Id1) (Fig. 2C). Tumor neurospheres (NS) expressing mutant ACVR1 (mACVR1 NS) also exhibited elevated levels of phospho-smad1/5 (Fig. 2D) and elevated levels of Id1 (Fig. 2D). To test the effect of a specific inhibitor of ACVR1, LDN-214117 on phospho-Smad1/5 signaling (Fig. 2E), we treated mACVR1 NS with increasing concentrations of LDN-214117. We observed decreased levels of phospho-Smad1/5 and Id2, while the levels of total Smad1 remain unchanged. Expression of ID1 was also confirmed on human DIPG cultures: SU-DIPG VI (wt-ACVR1) and SU-DIPG XX1 (mACVR1) (Supplementary Fig. S5).

### mACVR1 differentially regulates TGF-beta pathway and pathways related to stem cell maintenance and focal adhesion

RNAseq analysis of mACVR1 vs wt-ACVR1 NS identified genes that were differentially regulated (1.5-fold; FDR-corrected P < 0.05) (Fig. 3A). Gene ontology (GO) terms that were over represented in the set of differentially expressed genes (DE) included: developmental processes, cell migration, neuron differentiation, neurogenesis, nervous system development, cellular developmental process, cell development, cell proliferation, neuron development, cell surface receptor signaling pathway, neuron projection development, cell adhesion, positive regulation of developmental process, and positive regulation of cell differentiation (FDR correction; minimum 30 DE genes per term) (Fig. 3B). The top three pathways that were impacted by the mutation in ACVR1 were focal adhesion (FDR corrected, p=0.004), the TGF-beta signaling pathway (FDR corrected, p=0.004), and signaling pathways regulating pluripotency of stem cells (FDR corrected, p=0.009) (Fig. 3C). Over expressed genes within the TGF-Beta pathway included ACVR1, and inhibitor of DNA binding genes Id1, Id2, and Id3 (Fig. 3C). Gene set enrichment analysis (GSEA) suggests an enrichment in the response to BMP and the regulation of cell differentiation (Fig. 3D-F). Since one of the top signaling pathways was involved the regulation of pluripotency in stem cells, we evaluated whether mACVR1 tumors express CD133, CD44, and Aldh1, stem cell markers. Intracranial wt-ACVR1 or mACVR1 tumors were established in the pons of adult mice using SB derived NS. The results demonstrate that mACVR1 tumors have increased expression of the cancer stem cell markers CD133 (p=0.0002) and CD44 (p=0.0112) (Fig. 4A). We did not observe differences in the expression of another cancer stem cell marker, i.e., Aldehyde dehydrogenase 1 family, member A1 (Aldh1) (Fig. 4A). We next investigated the tumor initiating potential of wt-ACVR1 and mACVR1 NS *in vivo*. With wt-ACVR1 NS, the minimum number of cells required to generate brainstem gliomas with 100% penetrance was 1,000 cells, whereas, with mACVR1 NS it was possible to generate brainstem gliomas with 100% penetrance using 500 cells (Fig. 4B,C). These results indicate that mACVR1 NS have a greater tumor initiating potential.

**Figure 3.**
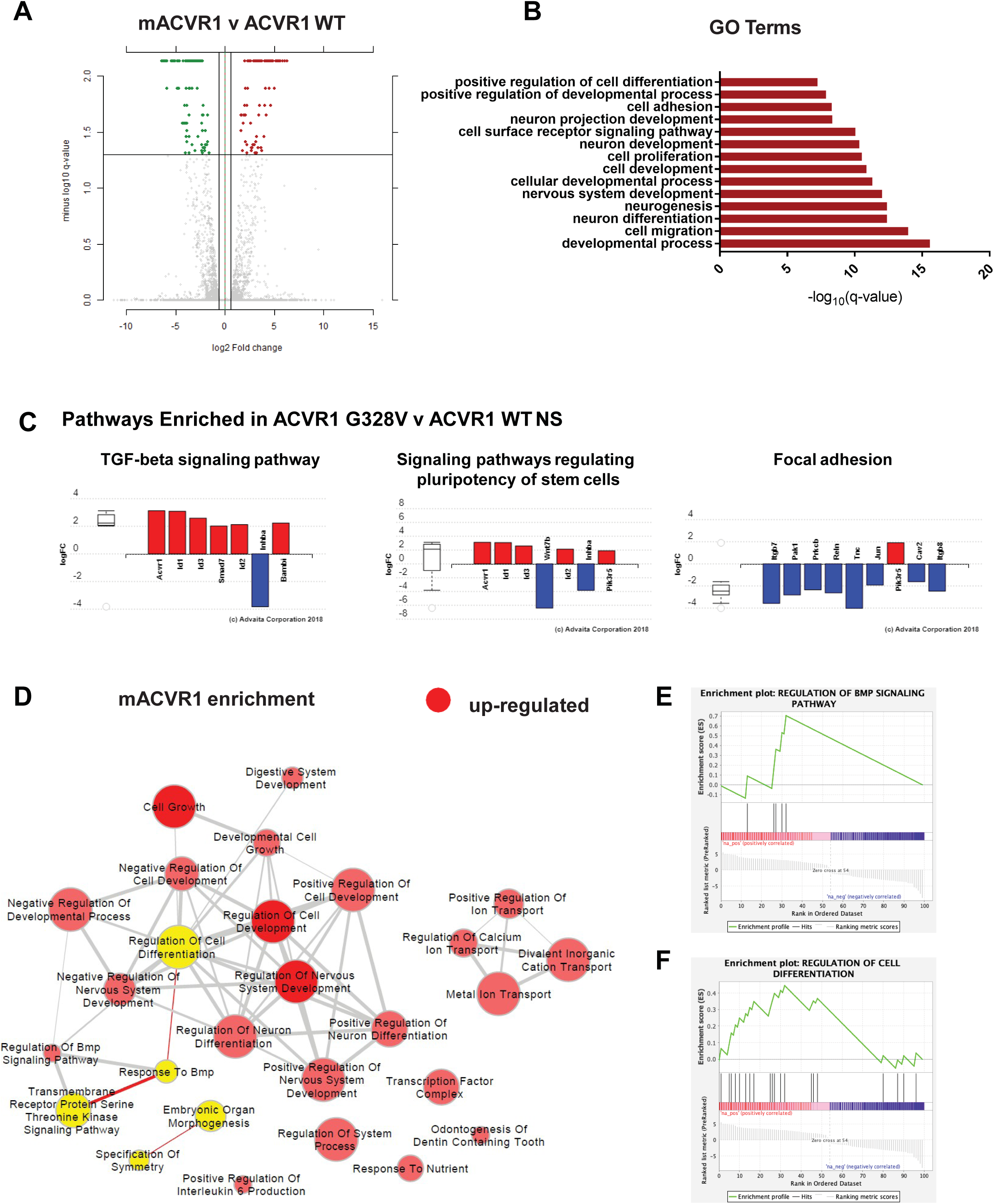
ACVR1^G328V^ upregulates genes involved in the TGF-ß /Smad signaling and pathways related to stem cell maintenance and focal adhesion. **(A)** Differential gene expression in mACVR1 tumors analyzed by RNA-seq. Volcano plot comparing differentially expressed genes in mACVR1 versus wt-ACVR1 mouse NS. The log 10 (FDR corrected p-values); q-values were plotted against the log 2 (Fold Change: FC) in gene expression. Genes upregulated by ≥ 1.3 fold and with a FDR corrected p-value < 0.05 are depicted as red dots; genes that were downregulated by ≥1.3 fold and with a FDR corrected p-value < 0.05 are depicted as green dots. **(B)** Gene Ontology (GO) enrichment analysis performed using iPathwayGuide for up- and down-regulated genes. The bar graph shows the GO biological process, selected for relevance to phenotype, that are differentially expressed between mACVR1 versus wt-ACVR1 NS. The –log10 q values of GO terms were plotted; FDR corrected P value = q value. **(C)** Differentially expressed pathway genes associated with mACVR1 in mACVR1 versus wt-ACVR1 mouse NS. Plot of the log (FC) of genes with an FDR corrected p-value < 0.05. **(D)** Pathway enrichment maps of DE genes in mACVR1 versus wt-ACVR1 NS. Clusters of nodes depicted in red illustrate differentially upregulated pathways resulting from GSEA (P < 0.05; FDR < 0.5). The yellow highlighted nodes indicate up-regulated GO terms containing ACVR1. **(E)** GSEA enrichment plot of regulation of BMP signaling pathway genes identified by RNA-Seq analysis in mACVR1 versus wt-ACVR1 mouse NS. **(F)** GSEA enrichment plot of regulation of cell differentiation genes identified by RNA-Seq analysis in mACVR1 versus wt-ACVR1 mouse NS.

**Figure 4.**
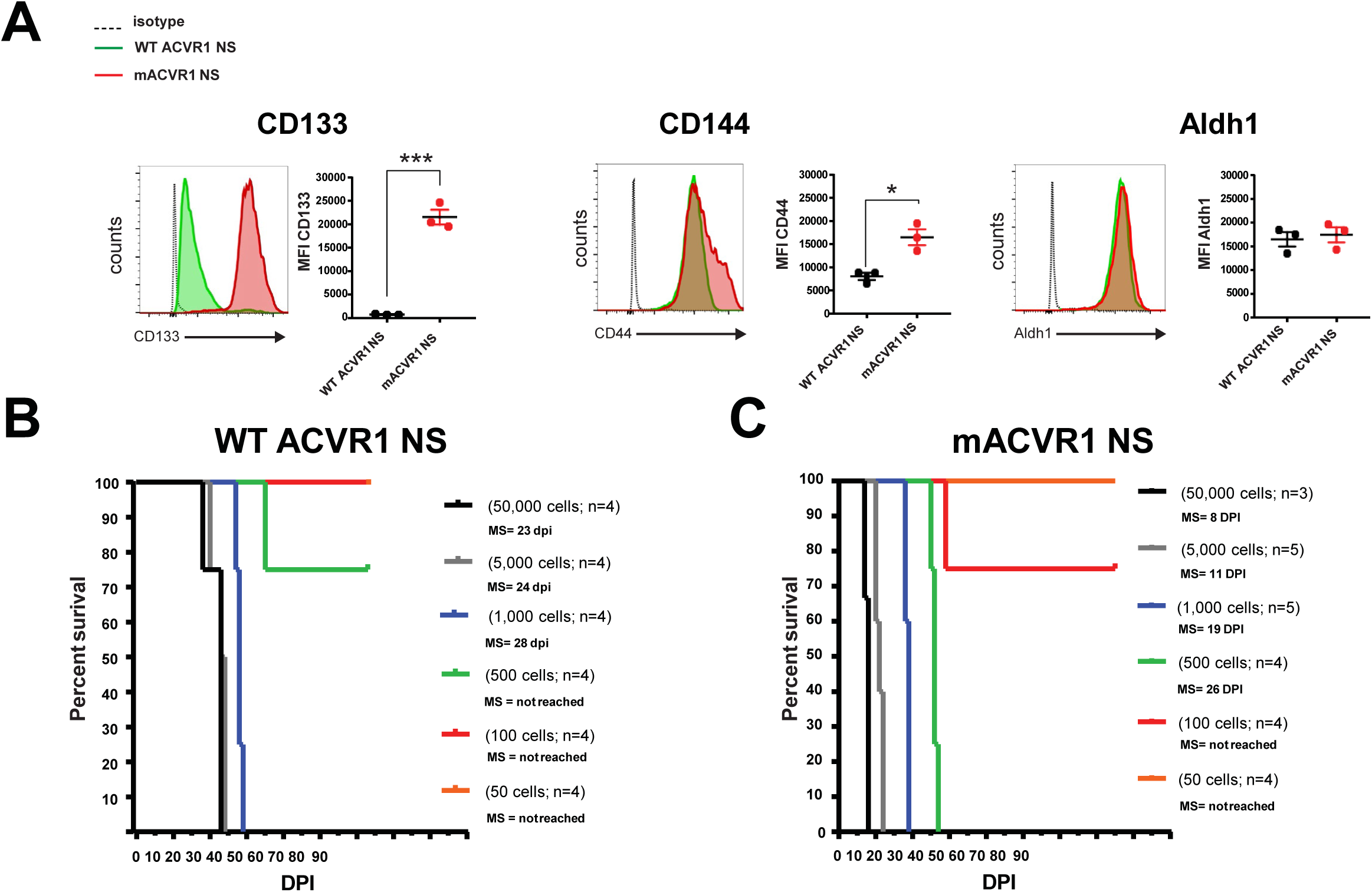
Increased expression of stem cell markers and greater tumor initiation potential in mACVR1 NS compared to WT ACVR1 NS. **(A)** Levels of CD133, CD44, and Aldh1 in the (tumor microenvironment) TME of mice bearing wt-ACVR1 or mACVR1 was assessed when animals displayed signs of tumor burden. Representative histograms display each marker’s expression (green = wt-ACVR1, red = mACVR1). MFI = mean fluorescence intensity; * p < 0.05; *** p < 0.001; unpaired *t* test. Bars represent mean ± standard error of the mean (SEM) (n = 3 biological replicates). (**B-C**) In vivo tumor initiation capacity. Kaplan-Meier survival curves for mice intracranially implanted with different numbers of wt-ACVR1 **(B)** or mACVR1 NS **(C)** as indicated in the plot. MS: median survival.

### Pre-clinical Testing of Immune Stimulatory Gene Therapy using SB derived NS

Due to the invasive nature of brainstem gliomas and their refractive response to current therapies we wanted to assess the efficacy of immune stimulatory TK/Flt3L gene therapy using an immune competent, intracranial mouse model of brainstem glioma. To generate this model, we utilized NS expressing mACVR1. The release of DAMPs is crucial for the success of TK/Flt3L mediated therapy, therefore, we first assessed whether mACVR1 NS released DAMPs *in vitro* after treatment with GCV alone, TK alone, or GCV+TK, with, or without 3 Gray (Gy) of irradiation (IR). We found that mACVR1 NS treated with GCV+TK released increased levels of calreticulin (p<0.0001; Fig. 5A), high mobility group box 1 protein (HMGB1) (p<0.0001; Fig. 5A), adenosine triphosphate (ATP) (p = 0.0071; Fig. 5A). We also observed that the combination of GCV+TK with IR further increased the release of calreticulin (p<0.0001; Fig. 5A), HMGB1 (p=0.0375; Fig. 5A), and ATP (p = 0.0158; Fig 5A) release by the mACVR1 NS.

**Figure 5.**
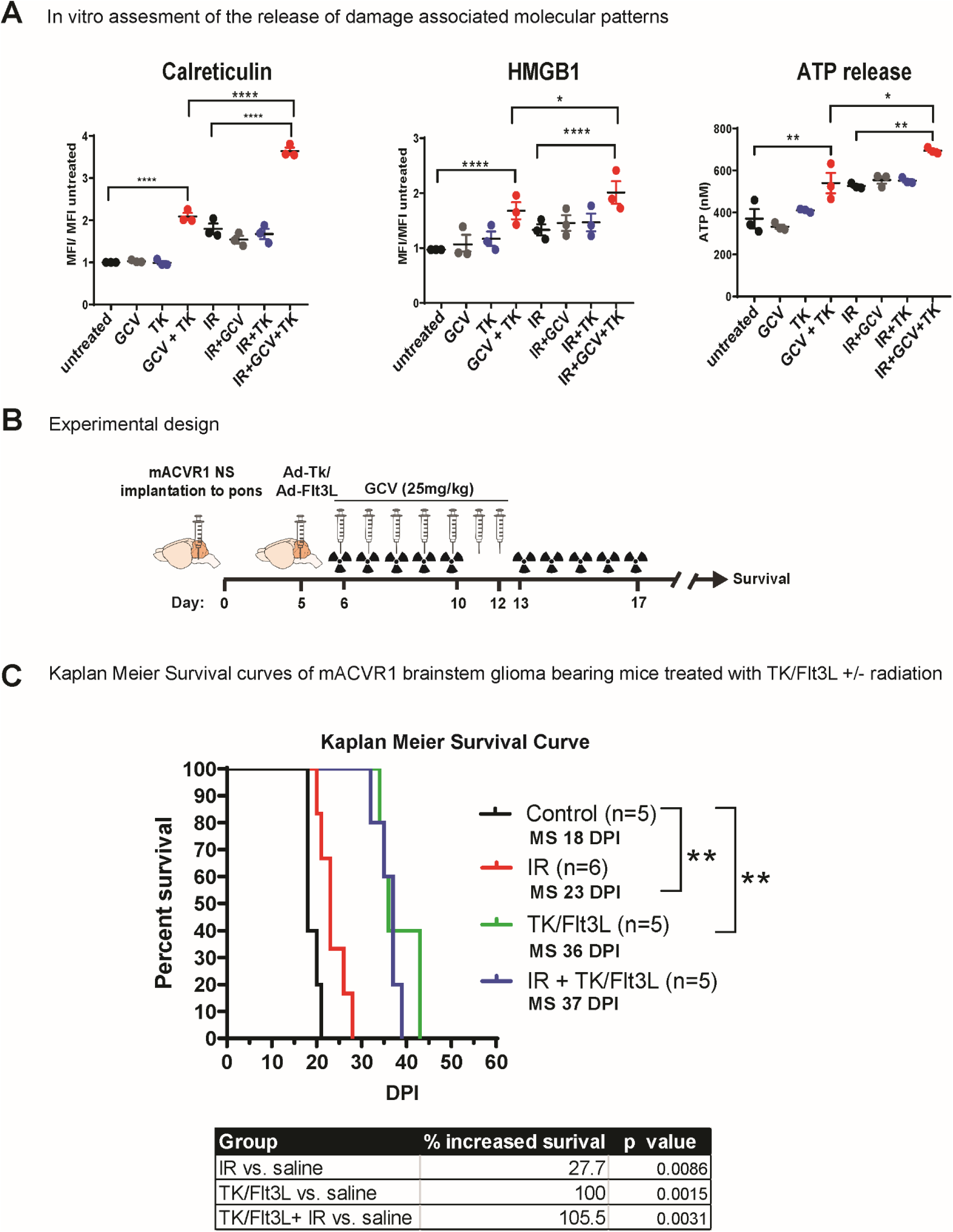
Efficacy of Immune-Stimulatory Gene Therapy. **(A)** wt-ACVR1 or mACVR1 NS were treated with TK (500 MOI) and/ or 3 Gray (Gy) ionizing radiation (IR), followed by GCV (25 µM). Levels of calreticulin or HMGB1 on tumor NS were assessed via flow cytometry analysis 24hrs post treatment. MFI = mean fluorescence intensity; * p < 0.05; **** p < 0.0001; two-way ANOVA, followed by Turkey’s multiple comparisons. Levels of ATP released into the media were assessed using a colorimetric assay. **(B)** Experimental design showing mice bearing mACVR1 brainstem gliomas treated with saline or TK/FLT3L gene therapy on day 5 post tumor implantation, followed by intraperitoneal administration of GCV (25 mg/kg) on day 6-12. Radiation was administered at a dose of 2Gy/Day for 5d for two weeks 5 days’ post tumor implantation; mice were monitored for survival. **(C)** Kaplan Meier survival analysis for mice bearing mACVR1 tumors treated with saline (n = 5), IR (n = 6), TK+ GCV (n=5), or IR+TK+GCV (n=5). Data were analyzed using log-rank (Mantel-Cox) test; ** p < 0.005.

Having established that mACVR1 NS release calreticulin, HMGB1, and ATP *in vitro* following treatment with either IR or TK+GCV, we next implanted mACVR1 NS into the pons of immune competent adult C57BL/6 mice to generate a transplantable brainstem model amenable to pre-clinical therapeutic implementation. Brainstem tumors derived using the transplantable model are positive for Ki67 (proliferating cells), GFAP (astrocyte marker), Vimentin (astrocytes and ependymal cells), Iba1 (microglia), but negative for MBP (a marker of mature oligodendrocytes) (Supplementary Fig. S6). The tumors are also positive for pERK1/2 and pMEK1/2 demonstrating activation of the RTK-RAS-PI3K signaling pathway (Supplementary Fig. S6). We also observed expression of Id1 a downstream target gene of BMP-Smad1/5 signaling (Supplementary Fig. S6). Five days’ post intracranial tumor implantation into the pons (Fig. 5B), mice were assigned to four treatment groups as indicated in Figure 5C. Our results demonstrate that Ad-TK/Ad-Flt3L therapy is more effective in prolonging the median survival (MS) of mACVR1 brainstem glioma compared to standard of care alone (MS=36 days post implantation (dpi) for TK/Flt3L group vs. 23 dpi for IR group; p = 0.0014, Mantel-Cox test) or vs. saline control (MS=18dpi; p = 0.0015, Mantel-Cox test) (Fig. 5C). Combining standard of care with gene therapy led to an improved median survival (MS=37 dpi), but did not significantly affect the efficacy of Ad-TK/Ad-Flt3L alone (Fig. 5C).

We next aimed to investigate whether Ad-TK/Ad-Flt3L treatment recruits anti-brainstem glioma specific T cell infiltration into the tumor immune microenvironment (TME). To do this, we used the surrogate tumor antigen ovoalbumin (OVA)(18,28). Using OVA-expressing mACVR1 cells (mACVR1-OVA NS), we were able to quantify tumor specific CD8 T cells in the TME through the use of the SIINFEKL-H2Kb tetramer (Fig. 6A, B). We observed a 3.8-fold increase in the frequency of tumor specific CD8 T cells in the TME after treatment with Ad-TK/Ad-Flt3L gene therapy (p = 0.0007; ***, Fig. 6B). To test the impact of Ad-TK/Ad-Flt3L gene therapy on the activation status of CD8 T cells in the TME, we stained effector T cells for the expression of IFNγ after re-stimulating CD8 T cells isolated from the TME with mACVR1-OVA NS-lysate for 24 hrs. Our data show that IFNγ is increased 6.7 fold (p < 0.0001; ****) in CD8 T cells from Ad-TK/Ad-Flt3L gene therapy treated mice compared to saline controls (Fig. 6C). To test if Ad-TK/Ad-Flt3L gene therapy affected antigen-specific T cell proliferation we labeled splenocytes with 5-(and 6)-carboxyfluorescein diacetate succinimidyl ester (CFSE), and stimulated them with the OVA cognate SIINFEKL peptide. As a positive control we also stimulated splenic T cells from OT-1 mice that have their TCR engineered to recognize the SIINKEKL peptide. Our results show that the percentage of T cells that proliferated in response to the SIINKEKL peptide was greater (1.7 fold, p < 0.0001) in mice treated with Ad-TK/Ad-Flt3L gene therapy compared to the saline treated control group (Fig. 6D). Additionally, the cytotoxicity of T cells isolated from the spleen of animals treated with Ad-TK/Ad-Flt3L gene therapy was observed to be significantly higher (1.8 fold, p < 0.0001 at 20:1 ratio) (Fig. 6E) when compared with the saline treated group indicating that gene therapy significantly enhanced the cytotoxic activity of splenic T cells.

**Figure 6.**
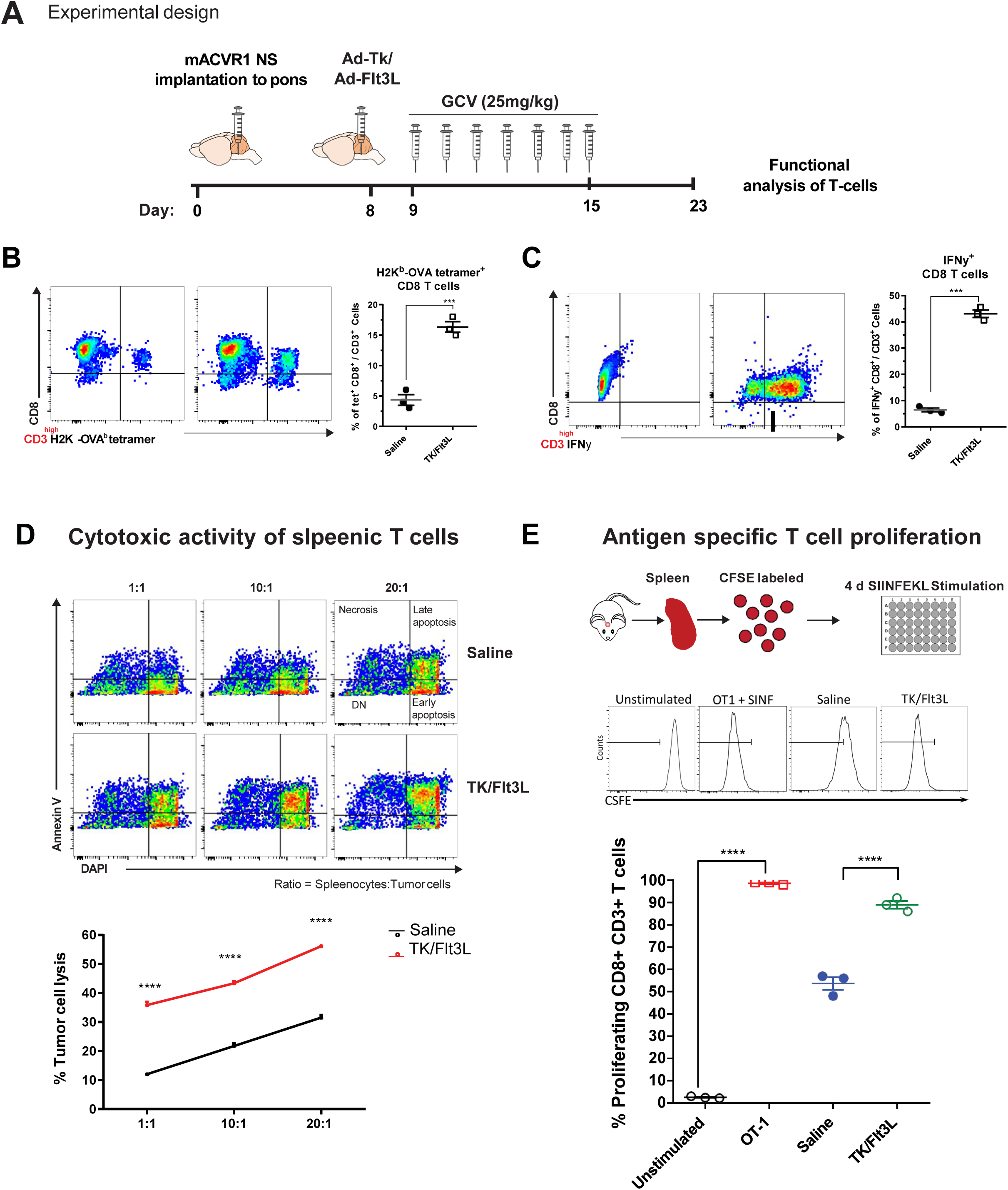
(A) Immune stimulatory gene therapy induces tumor specific T-cell infiltration. Experimental design showing mice bearing mACVR1-OVA brainstem gliomas treated with saline or TK/FLT3L gene therapy on day 8 post implantation, followed by intraperitoneal administration of GCV (25 mg/kg) on day 9-15. **(B)** Tumor specific CD8 T cells within the TME of mACVR1-OVA tumors were analyzed by staining for SIINFEKL-K^b^ tetramer. **(C)** Activation status of CD8 T cells within the TME was analyzed by staining for IFNγ after stimulation with tumor lysate. Representative flow plots for each group are displayed; *** p < 0.001, **** p < 0.0001; unpaired t test. Bars represent mean ± SEM (n = 3 biological replicates, where resected tumors from 3 animals where pooled for the control group and 5 animals were pooled for the gene therapy group). **(D)** Splenocytes from saline or TK/Flt3L treated mice were incubated with mACVR1 NS for 24 hours at the indicated ratios. Lysis of tumor cells was assessed by using Annexin-V and DAPI. mACVR1 NS undergoing late apoptosis were identified as Annexin-V+/DAPI+ cells. Data were compared using unpaired t test; **** p < 0.0001. Bars represent mean ± SEM (n = 3 biological replicates). **(E)** Experimental design showing splenocytes from saline or TK/FLT3L treated mACVR1-OVA tumor bearing mice labeled with CFSE and then stimulated with 100 nM of SIINFEEKL peptide for four days in culture to assess CD8+ T cell proliferation. Histograms show representative CFSE staining from unstimulated splenocytes (negative control), OT-1 splenocytes undergoing rapid proliferation in response to SIINFEKL (positive control), and the effect of SIINFEKL-induced T cell proliferation on splenocytes from saline or TK/Flt3L treated mACVR1-OVA tumor bearing mice. Quantification of splenocytes undergoing T cell proliferation; **** p < 0.0001; one-way ANOVA followed by followed by Turkey’s multiple comparisons. Bars represent mean ± SEM (n = 3 biological replicates).

To evaluate any potential adverse effects of delivering gene therapy into the brainstem we performed a detailed histopathological analysis of brains from non-tumor bearing animals. After the animals were treated intracranially with Ad-TK/Ad-Flt3L, GCV was administered intraperitoneally one day after adenoviral injection for seven days. Brains were harvested for neuropathology analysis 24 hours after the last dose of GCV was administered. Architectural integrity was assessed by H&E staining and by immunohistochemistry using GFAP (astrocytes), MBP (myelin sheaths, oligodendrocytes) and Iba1 (microglia). No gross tissue abnormalities were observed in response to TK/Flt3L therapy compared to the saline controls (Supplementary Fig. S7). There was also no change in GFAP or MBP expression in response to TK/Flt3L therapy indicating that the brain architecture was unaffected (Supplementary Fig. S7). There was no increase in Iba1 expression in the animals treated with TK/Flt3L therapy indicating that gene therapy does not induce inflammation in normal brain tissue 8 days post treatement (Supplementary Fig. S7).

To assess potential systemic toxicity due to TK/Flt3L or IR treatment, we performed H&E staining on liver sections and complete hematological and serum biochemical analysis in tumor bearing animals from the saline, TK/Flt3L, and TK/Flt3L + IR treatment groups. The liver sections from all treatment groups had normal hepatocyte architecture and did not show signs of inflammation or necrosis (Supplementary Fig. S9). The white blood cell counts were within normal range for the saline, TK/Flt3L, and TK/Flt3L + IR groups, but significantly decreased in the IR group in comparison to the saline treated group (p < 0.0001, Supplementary Fig. S9). This is consistent with a study that found that whole brain radiation significantly decreased total white blood cell counts (29). Red blood cell, hemoglobin, hematocrit, platelet, lymphocyte, neutrophil, and monocyte counts were not significantly affected by TK/Flt3L or IR therapy (Supplementary Fig. S9). We did not find any significant changes in important enzymes involved in liver (ALT, AST) and kidney (BUN) function as a result of TK /Flt3L or IR therapies (Supplementary Fig. S9).

## Discussion

DIPG remains an incurable tumor with a poor prognosis (30). The tumors originate in the pons and infiltrate into sensitive regions of the brainstem precluding surgical resection (31). DIPGs are also resistant to radiation and chemotherapy (32). DIPG studies involving the development and implementation of novel therapies have utilized patient derived xenograft models, where human DIPG cells are implanted into the brain of immune deficient mice or rats to establish intracranial tumors (33,34). One limitation with those models is that immune suppressed animals cannot be used to test immunotherapies or perform immune-related mechanistic studies. In this study, we utilized the SB transposase system to generate an endogenous mouse model of mutant ACVR1 brainstem glioma. ACVR1 is frequently mutated to ACVR1 G328V in DIPG (4-7). The SB system efficiently and reproducibly integrates plasmid DNA into the neural progenitor cells’ host chromosomal DNA of neonatal mice, allowing for the functional assessment of the role of candidate DIPG genes in promoting tumor progression (19-22). Mutations in ACVR1 confer ligand independent activation of ACVR1, which is a mediator of the BMP signaling pathway (4,5,35,36). We show that SB tumors are localized in the brainstem and have increased downstream signaling of BMP as demonstrated by increased phospho-smad1/5 levels and Id1 (Fig. 2B, C; **Supplementary Fig. S5**).

Neurospheres (NS) harboring mACVR1 were derived from sleeping beauty generated tumors; they have stem cell-like properties and are grown in suspension as 3D cultures. Transcriptional profiling indicated an enrichment in the BMP signaling pathway in mACVR1 NS compared to WT ACVR1 tumor NS (Fig. 3D). These NS were then implanted into the pons of adult immune competent mice, enabling the generation of transplantable DIPG models and amenable for preclinical therapeutic studies. Currently, the treatment option for children with DIPG is limited to radiation to provide palliative care (1). Although radiation can temporarily provide symptomatic relief and extend survival by a few months, it can cause detrimental effects on the developing brain (37,38). Therefore, there is a dire need for new therapeutic interventions. This research focuses on the response of mACVR1 brainstem glioma tumors to immunotherapies.

There has been significant progress in the field of cancer immunotherapy leading to improvements in overall survival in many types of solid tumors (11-13,39,40). Advances in immunotherapies include the development of immune therapies targeting immune checkpoints, vaccine approaches against tumor antigens or dendritic cell vaccines designed to stimulate the adaptive immune response, adoptive cell therapy, oncolytic viral therapy, and immune stimulatory gene therapy (40-42). This has led to a growing number of clinical trials testing immunotherapies in DIPG (42).

In recent years, it has been established that activated immune cells are able to migrate and enter into the brain parenchyma, and that the brain has a functional lymphatic system that allows for transport of CNS-antigens to the draining lymph nodes (43,44). Thus, the brain is capable of mounting T cell-mediated adaptive immune responses, but, since there are very low numbers of local professional antigen presenting cells within the normal brain, it is not possible to prime a potent immune response against antigens localized in the brain parenchyma (45,46).

In adult glioblastoma (GBM) there is evidence of immune cell infiltration, but an immunosuppressive environment precludes effective anti-tumor immunity (41,47,48). GBM tumors establish an immunosuppressive environment by the release of immunosuppressive cytokines, such as TGF-β and IL-10, by the recruitment or induction of immunosuppressive cells, such as regulatory T cells, myeloid-derived suppressor cells, or tumor associated macrophages, and by the expression of immune checkpoint receptor ligands (47-53). In comparison to adult GBM, initial studies of the tumor microenvironment in DIPG have found that there is a low number of immune infiltrates in human DIPG tumors and that they do not express inflammatory cytokines and chemokines (54,55). These data provide support for the use of an immune modulatory therapeutic strategy to enhance the recruitment of immune cells into the tumor, with the aim of mounting an effective anti-DIPG immune response.

We have previously demonstrated that combined immune stimulatory gene therapy mediated through the delivery of adenoviruses encoding herpes simplex virus type 1 thymidine kinase (TK) and fms-like tyrosine kinase 3 ligand (Flt3L) leads to tumor regression and long term survival in several rodent models of glioblastoma(15,16,56-58). This therapy is based on inducing tumor cell death though expression of suicide gene TK (59,60). Tumor antigens and damage-associated molecular pattern molecules (DAMPs), such as calreticulin, high mobility group box1 (HMGB1), and ATP, are released by dying tumor cells (17,61). The effectiveness of this combination therapy also relies on Flt3L to recruit dendritic cells into the tumor microenvironment, while the release of HMGB1 stimulates TLR2-dependent activation of dendritic cells (15,60,62). Activated dendritic cells can then transport antigens to the draining lymph nodes and induce tumor specific T cell responses (15,60). Initial results from the first in human Phase 1 clinical trial of combined adenoviral delivery of TK and Flt3L for the treatment of adult GBM are promising and report that the therapy was well tolerated (63). However, it has been established that the biology of DIPG and GBM is different (64). Therefore, it is necessary to assess the safety and efficacy of TK/Flt3L therapy in pre-clinical models of brainstem glioma. Herein, we demonstrate that treatment with TK/Flt3L gene therapy in mice bearing mACVR1 brainstem gliomas stimulates a strong anti-tumor cytotoxic immune response leading to a significant increase in survival (Fig. 5C). Treatment with TK/Flt3L increased the frequency of tumor-specific CD8 T cells in the tumor microenvironment and increased toxicity as demonstrated by enhanced IFNγ production (Fig. 5 E, F).

Additionally, delivery of TK/Flt3L into the normal brainstem did not induce any local or systemic cytotoxicity. Hematoxylin and eosin staining and immunostaining for GFAP (astoryctes), MBP (myelin sheaths, oligodendrocytes), and Iba1 (activated macrophages and microglia) was used to assess local toxicity, and we observed no architectural abnormalities or overt inflammation as a result of TK/Flt3L therapy. Histological examination of hematoxylin and eosin stained liver tissue did not show signs of inflammation, necrosis, or alterations in normal hepatocyte structure. Hematological toxicity, assessed by complete blood count and serum chemistry analysis, indicated that TK/Flt3L therapy did not induce any toxicity as values from a TK/Flt3L treated group were not significantly altered when compared to saline treated animals. Our results are consistent with other pre-clinical studies that report the brainstem can tolerate adenoviral mediated immunotherapy (65,66). Results from a clinical trial (NCT03178032) utilizing adenoviral vector delivery of an oncolytic virus into the pons of DIPG patients will also shed light on the feasibility and toxicity of intratumoral adenoviral delivery into the brainstem (67).

In conclusion, we demonstrate the amenability of the SB transposase to be used to develop genetically and histologically accurate models of DIPG expressing mutant ACVR1 and provide compelling evidence that warrants further development of conditionally cytotoxic immune stimulatory gene therapy for the treatment of DIPG. We anticipate that in the clinic, this approach could also be used in combination with immune checkpoint blockade to further enhance the therapeutic efficacy of the TK/Flt3L-mediated anti-brain stem glioma immune response.

## Acknowledgments

We thank J. Ohlfest (University of Minnesota, deceased) for providing the SB model plasmids and Jeff Wrana for providing the ACVR1 WT plasmid. We thank the support and academic leadership of Dr. Karin Muraszko, and the administrative support of Angela Collada. The content is solely the responsibility of the authors and does not necessarily represent the official views of the NIH.

