## Supplementary Materials for "Therapeutic efficacy of an immune stimulatory gene therapy strategy in a mouse model of high grade brainstem glioma"

**This PDF file includes:**

Supplementary Figures S1-S9

Supplementary Table 1

Supplementary Methods

References of Supplementary Methods

### Supplementary Fig S1

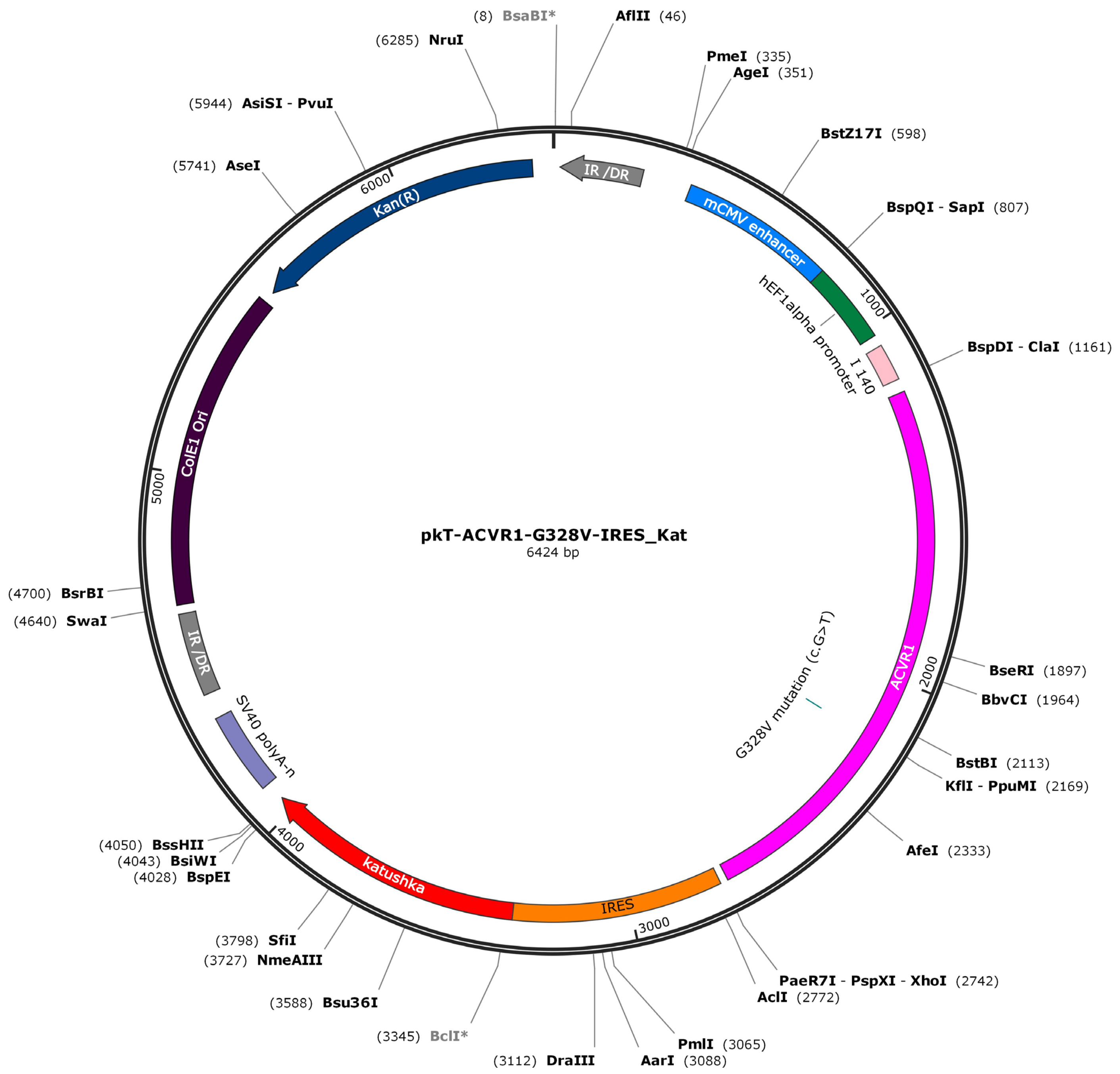

**Supplementary Figure S1:** Plasmid encoding mACVR1 G328V was cloned in between IR/DR sites and it is recognized by Sleeping Beauty.

### Supplementary Fig S2

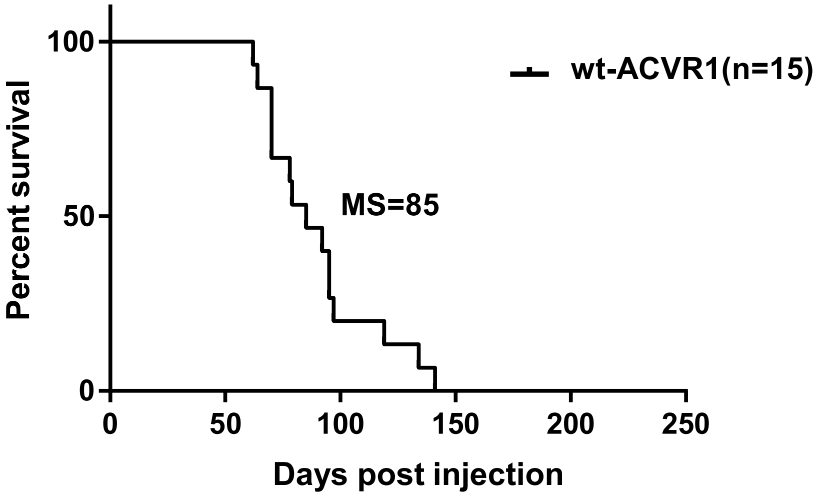

**Supplementary Figure S2**: Kaplan Meier survival curve for genetically engineered mice harboring wt-ACVR1 tumors. MS = median survival.

#### Supplementary Fig S3

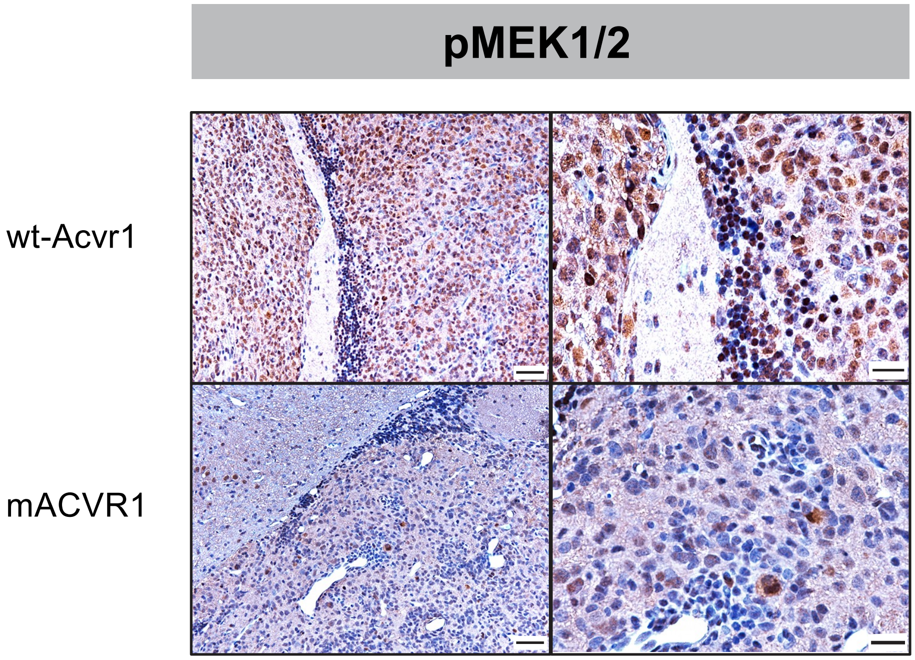

**Supplementary Figure S3:** Immunohistochemistry (IHC) staining for pMEK1/2 in paraffin embedded SB brainstem tumor sections (wt-ACVR1 and mACVR1). Scale bar on the left panel is 50 µm, scale bar for the right panel is 20 µm.

#### Supplementary Fig S4

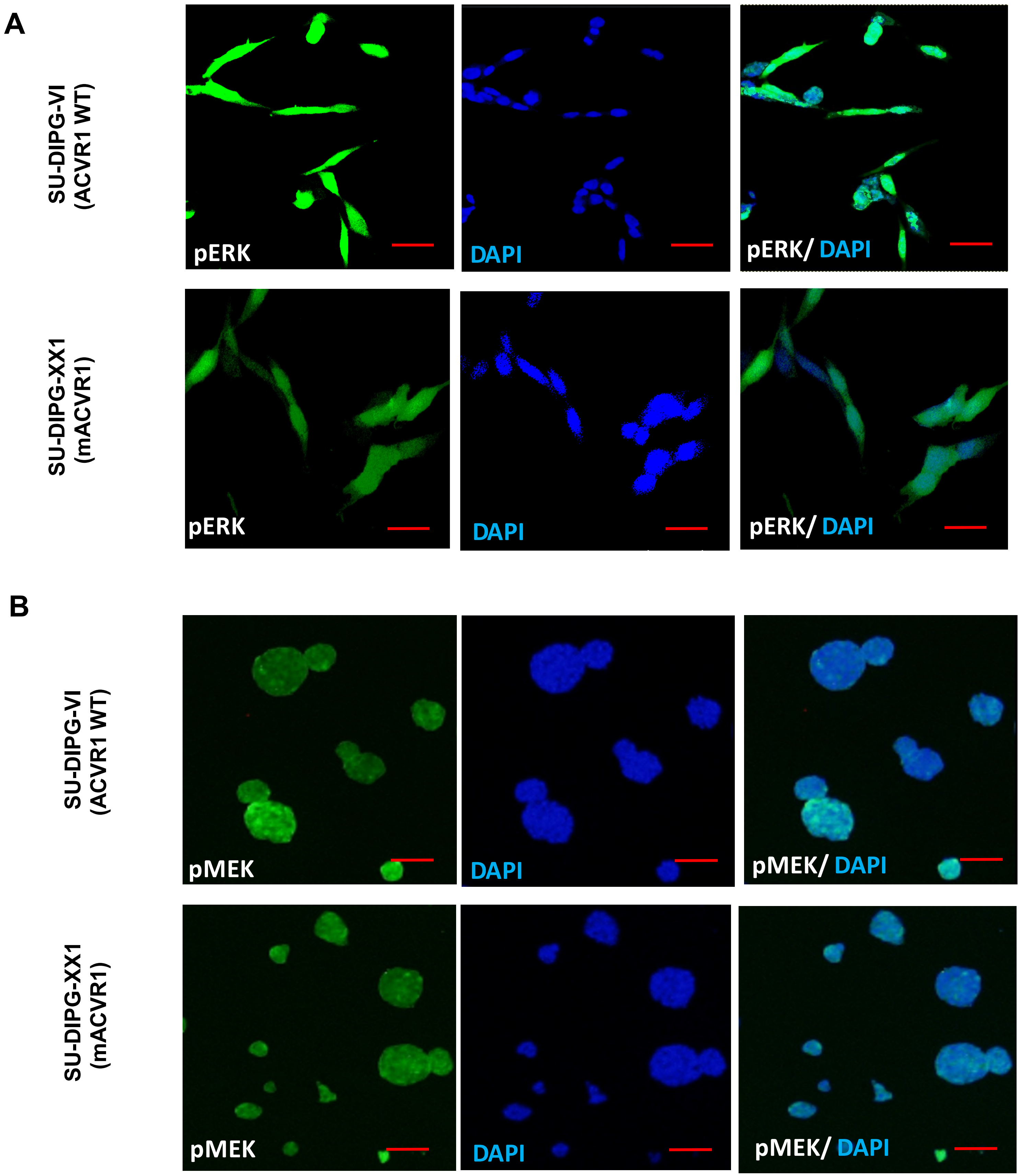

**Supplementary Figure S4**: Immunocytochemistry (ICC) staining for pERK1/2 and pMEK1/2 on human DIPG cells: SU-DIPG VI (wt-ACVR1) and SU DIPG XX1 (mACVR1). Scale bar = 50 μm.

#### Supplementary Fig S5

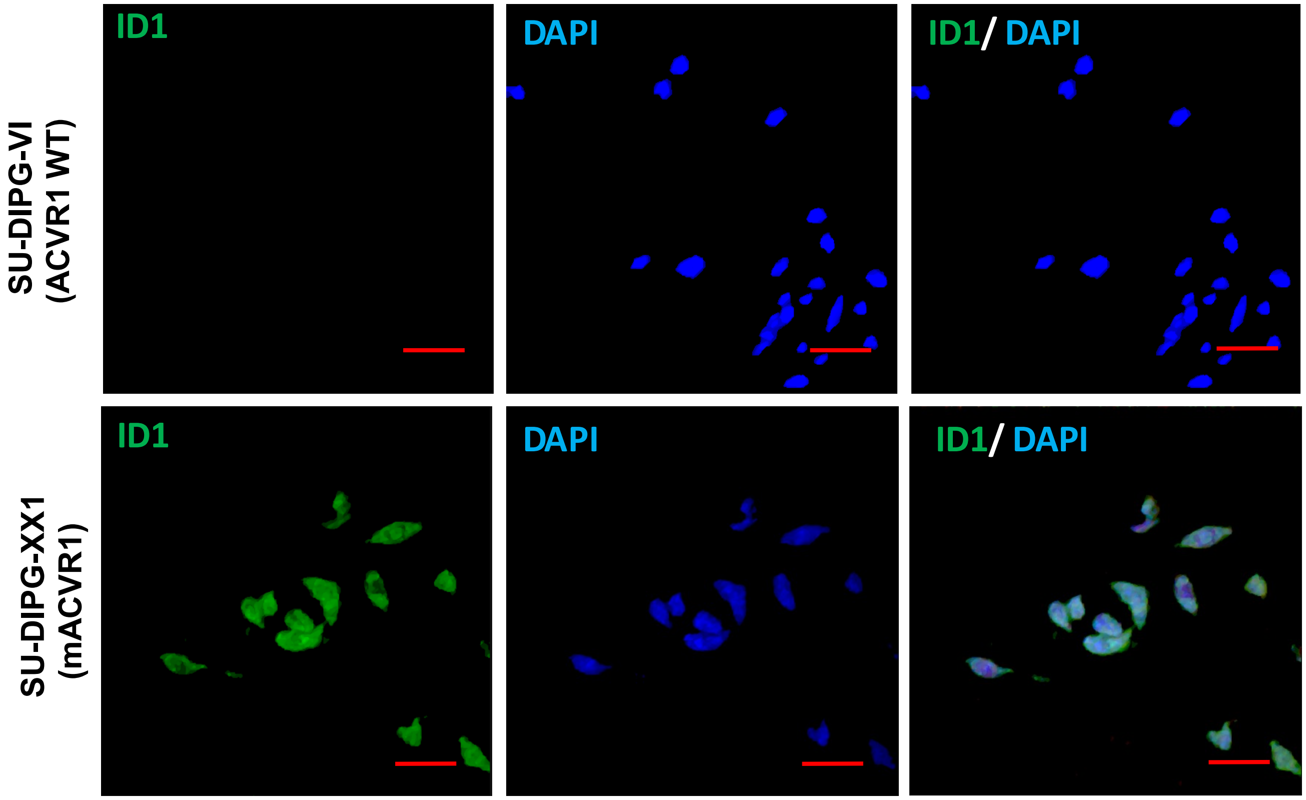

**Supplementary Figure S5**: Immunocytochemistry (ICC) staining for Id1 on human DIPG cells: SU-DIPG VI (wt-ACVR1) and SU DIPG XX1 (mACVR1). Scale bar = 50 μm.

### Supplementary Figure S6

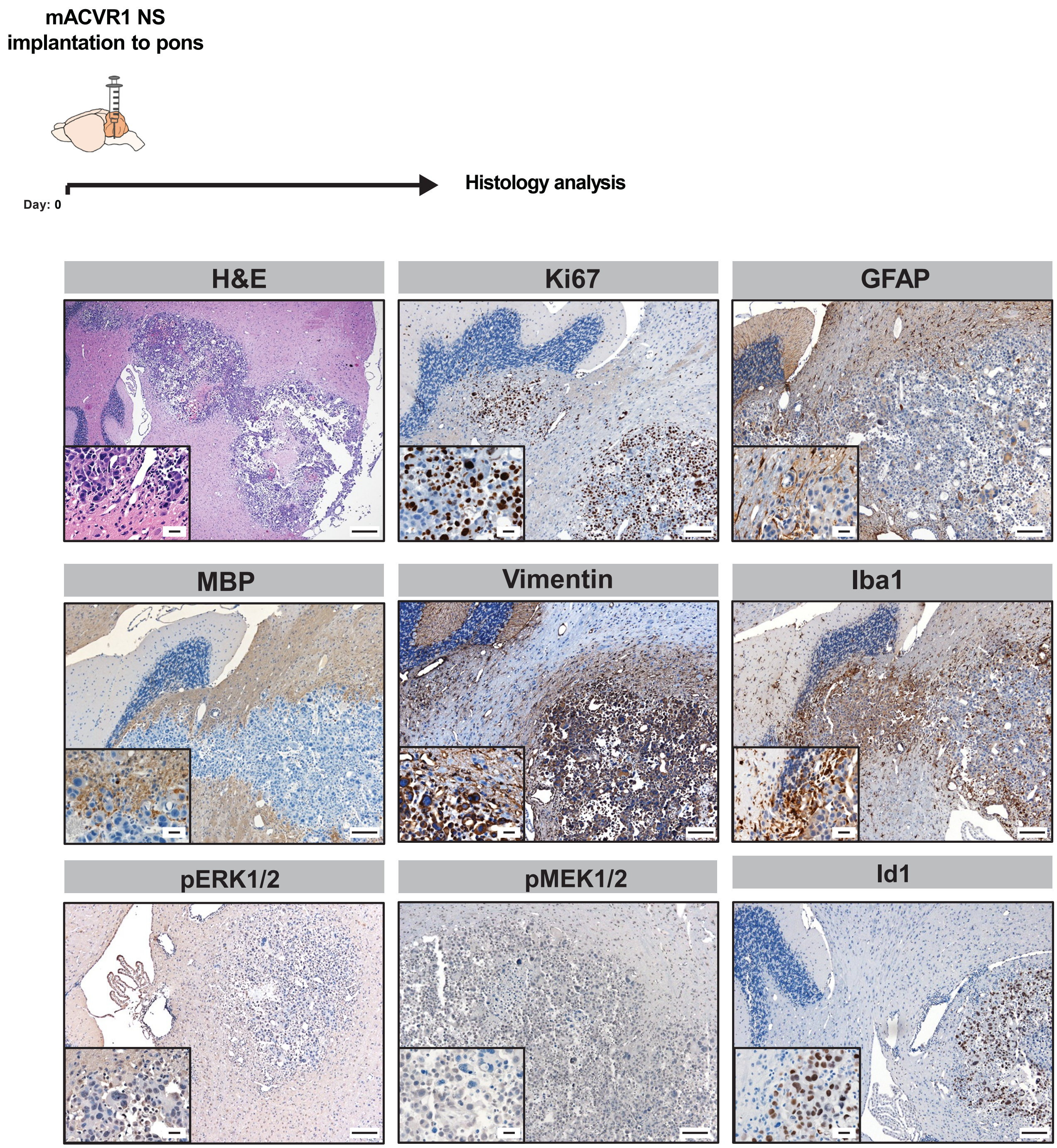

**Supplementary Figure S6**: Mice bearing mACVR1 intracranial brainstem gliomas were processed for histological analysis when they displayed signs of tumor burden. Paraffin embedded mACVR1 tumor sections were stained with Hematoxylin and Eosin (H&E), Ki67 (proliferating cells), GFAP (astrocytes), MBP (myelin sheaths and oligodendrocytes), Vimentin (ependymal cells and astrocytes), Iba1 (microglia), pERK1/2, pMEK1/2, or Id1. Scale bar = 200 μm for the H&E. Scale bar = 100 μm for the immunostains. Scale bar for all magnified insets = 20 μm.

#### Supplementary Fig S7

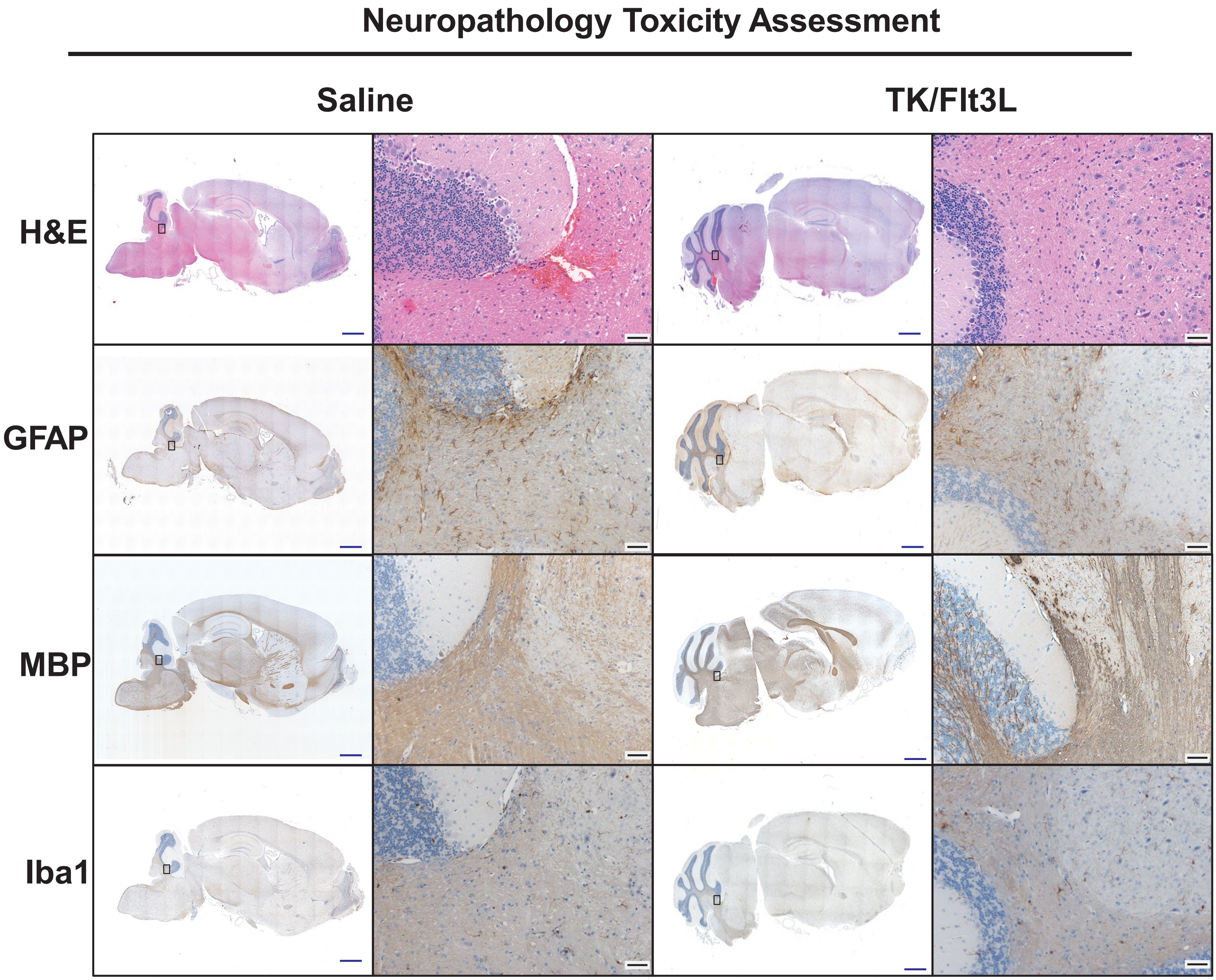

**Supplementary Figure S7**: Local toxicity of adenoviral delivery of TK/Flt3L into normal brainstem. Mice were injected in the pons with Ad-TK and Ad-Flt3L, or saline. GCV was administered intraperitoneally 24 h post vector delivery for 7 days. 24 h after the last dose of GCV was administered neuropathological analysis of the brain was assessed by H&E staining and immunostaining for GFAP (astrocytes), MBP (myelin sheaths, oligodendrocytes), and Iba1 (microglia). Black box represents magnified area. Blue scale bar = 1 mm. Black scale bar = 50 µm.

### Supplementary Fig S8

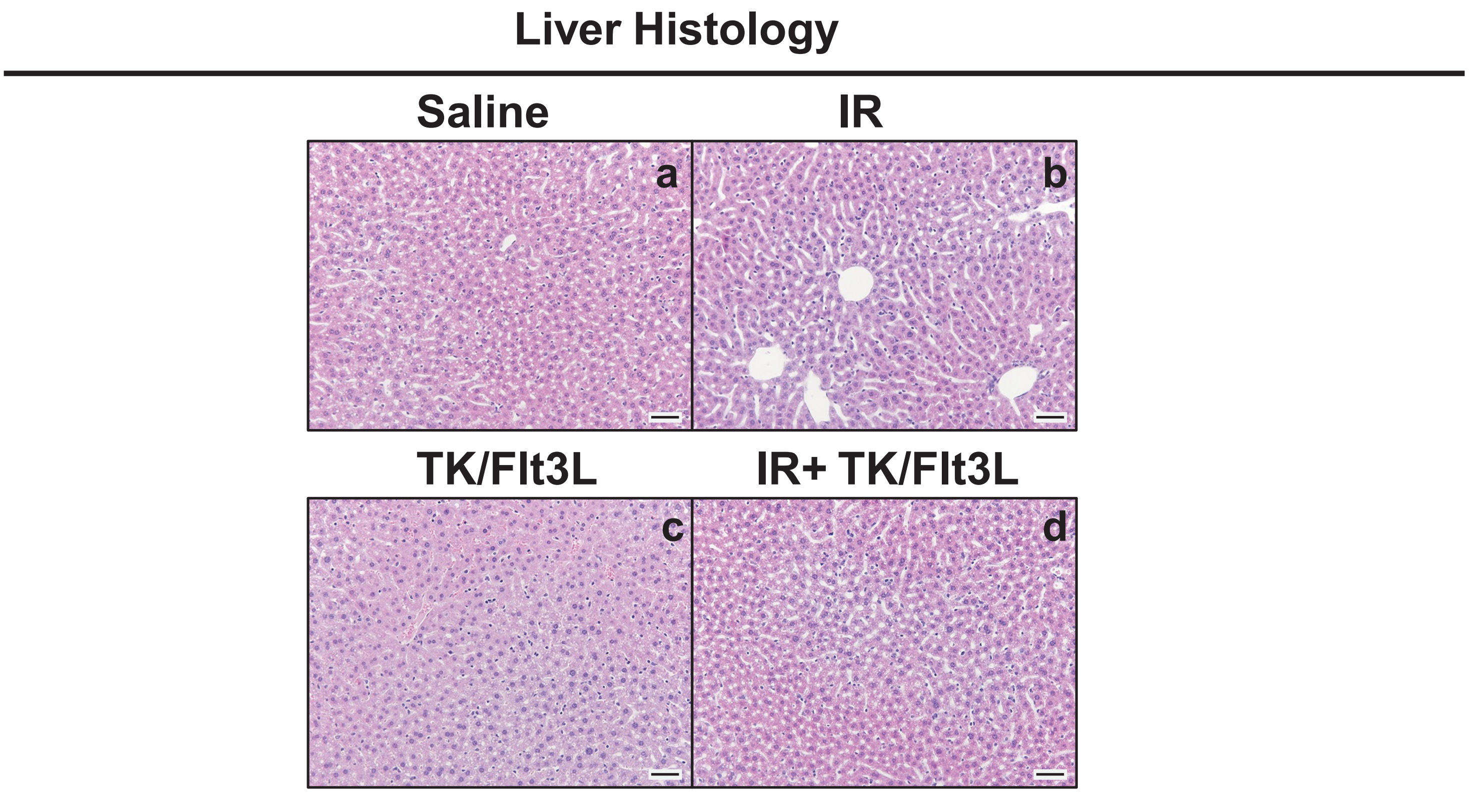

**Supplementary Figure S8**: Mice bearing mACVR1 brainstem gliomas were treated with saline or TK/FLT3L gene therapy on day 5 post tumor implantation, followed by intraperitoneal administration of GCV (25 mg/kg) on days 6-12. Radiation was administered at a dose of 2 Gy/Day for 5d for two weeks 5 days' post tumor implantation. Livers were processed for histology on day 23. Bright field images of paraffin embedded liver sections stained with H&E of animals that were treated with (a) saline, (b) IR, (c) Tk/Flt3L + GCV or (d) Tk/Flt3L + GCV+ IR. Scale bar = 50 µm.

#### Supplementary Fig S9

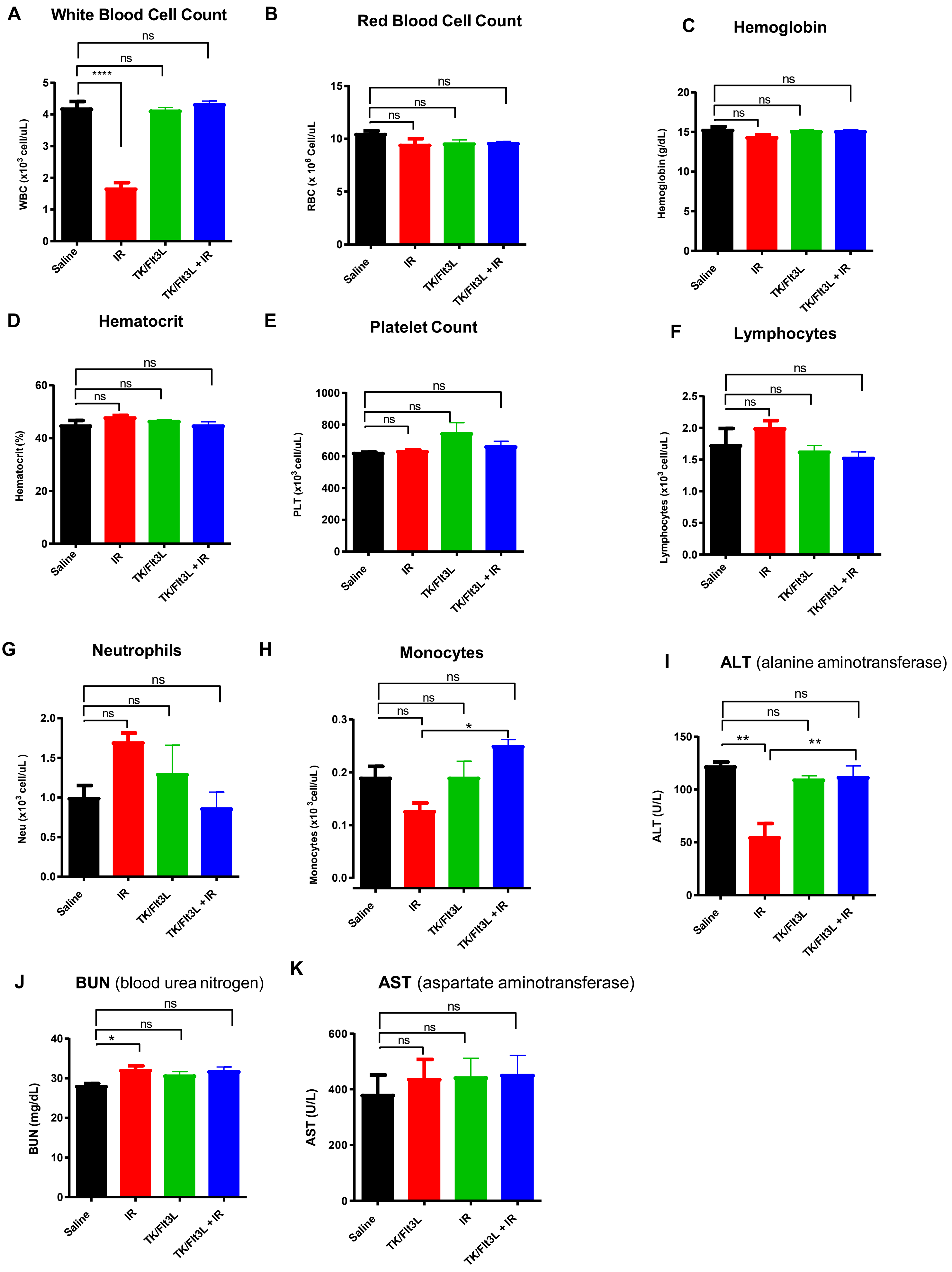

**Supplementary Figure S9:** Hematology and serum chemistry. Mice bearing mACVR1 brainstem gliomas were treated with saline or TK/FLT3L gene therapy on day 5 post tumor implantation, followed by intraperitoneal administration of GCV (25 mg/kg) on days 6-12. Radiation was administered at a dose of 2 Gy/Day for 5d for two weeks 5 days' post tumor implantation. Blood and serum samples were drawn on day 23 to assess clinical parameters. **(A)** White blood cell (WBC), **(B)** red blood cell (RBC), **(C)** hemoglobin, **(D)** hematocrit, **(E)** Platelet (PLT), **(F)** Lymphocytes, **(G)** Neurotrophils (Neu), **(H)** Monocytes, **(I)** alanine aminotransferase (ALT), **(J)** blood urea nitrogen (BUN), **(K)** aspartate aminotransferase (AST).

#### Supplementary Table 1. Antibodies

| Analysis | Antibody | Company | Catalog number | Dilution |
|---|---|---|---|---|
| IHC-DAB | Rabbit polyclonal anti-OLIG2 | Millipore Sigma | AB9610 | 1:200 |
| IHC-DAB | Rabbit polyclonal anti-GFAP | Millipore Sigma | AB5804 | 1:1000 |
| IHC-DAB | Chicken polyclonal nestin | Novus Biologicals | NB100-1604 | 1:100 |
| IHC-DAB | Rabbit polyclonal anti-SOX2 | Abcam | ab97959 | 1:500 |
| IHC-DAB | Rabbit Anti-Ki67 | Abcam | ab15580 | 1:100 |
| IHC-DAB | Rabbit monoclonal anti-p-ERK1/2 | Cell Signaling | 4370 | 1:100 |
| IHC-DAB | Rabbit polyclonal anti p-MEK-1/2 | Santa Cruz Biotechnology | sc-7995-R | 1:100 |
| WB | Rabbit anti phospho Smad1/Smad5/Smad8 (Ser463/465) | Millipore Sigma | AB3848-I | 1:500 |
| IHC Fluorescence | Rabbit anti-phospho Smad1/Smad5/Smad8 (Ser463/465) | Millipore Sigma | AB3848-I | 1:50 |
| IHC DAB | Rabbit Monoclonal Anti-Mouse Id1 antibody | Biocheck | BCH-1/37-2 | 1:100 |
| WB | Rabbit Monoclonal Anti-Mouse Id1 antibody | Biocheck | BCH-1/37-2 | 1:500 |
| WB | Rabbit Monocolonal Anti-Id2 | Calbioreagents | 9-2-8 | 1:500 |
| WB | Mouse monoclonal anti-β-actin | Millipore Sigma | A1978 | 1:10,000 |
| WB | Rabbit polyclonal anti-Smad1 | Abcam | ab63356 | 1:500 |
| WB | Goat anti-rabbit | Dako | P0448 | 1:4000 |
| WB | rabbit anti-mouse | Dako | P0260 | 1:20,000 |
| Flow cytometry | Mouse monoclonal anti CD16/32 | Biolegend | 101335 | 1:200 |
| Flow cytometry | Mouse monoclonal anti CD45 | Biolegend | 103125 | 1:200 |
| Flow cytometry | Mouse monoclonal anti CD8 | Biolgened | 100711 | 1:200 |
| Flow cytometry | Mouse monoclonal anti CD3 | Biolegend | 100333 | 1:200 |
| Flow cytometry | SIINFEKL-H2Kb-tetramer-PE | MBL International | TB-5001-1 | 1:200 |
| Flow cytometry | IFNγ | Biolegend | 113603 | 1:200 |
| Flow cytometry | CD133 | Biolegend | 141203 | 1:200 |
| Flow cytometry | CD44 | Biolegend | 103005 | 1:200 |
| Flow cytometry | Aldh1 | R&D Systems | AAF5869-SP | 1:200 |
| IHC-DAB | Vimentin | Abcam | ab92547 | 1:500 |
| IHC-DAB | IBA1 | Abcam | ab178846 | 1:1000 |
| IHC-DAB | CD8 | Cederlane | 361003(SY) | 1:2000 |

### Supplementary Methods

#### Experimental model

All animal studies were conducted according to guidelines approved by the Institutional Animal Care and Use Committee at the University of Michigan. Animals were housed in an AAALAC accredited animal facility and had constant access to food and water; they were monitored daily for tumor burden. Males and females were used. The strain of mice utilized in the study was C57BL/6 (Jackson Laboratory, 000664).

A murine model of brainstem glioma was generated by employing the Sleeping Beauty (SB) transposon system to integrate plasmid DNA into the genome of postnatal day 1 (P1) mouse pups. The plasmids utilized were as follows: (i) SB Transposase and luciferase (pT2C-LucPGK-SB100X, henceforth referred to as SB/Luc) (ii) a short-hairpin against p53 (pT2-shp53-GFP4, henceforth referred to as shp53) or shp53-NO-GFP (iii) a constitutively active mutant of NRAS (pT2CAG-NRASV12, henceforth referred to as NRAS) with or without (4) mutant ACVR1 G328V (pkt-ACVR1-G328V-IRES-Katushka; henceforth referred to as mACVR1)(1-3). To create the mACVR1 plasmid we cloned pCMV5-ALK2-WT into pKT2-IRES-Katushka by blunt cloning. Then, we used the QuikChange II Site-Directed Mutagenesis Kit (Agilent, 200523) to introduce (c.983G>T, p.Gly328Val) mutation into pKT2-ALK2-WT-IRES-Katushka, to generate pKT2-ACVR1-G328V-IRES-Katushka (Addgene plasmid #77437). The resultant mACVR1 plasmid was confirmed by Sanger sequencing. SB/Luc, shp53 and NRAS plasmids were the generous gift of Dr. John Ohlfest (University of Minnesota, now deceased). The pCMV5-ALK2-WT plasmid was a generous gift from Jeff Wrana (Addgene plasmid #11741). All experiments were performed using post-natal day 1 (P1) or P2 wild-type C57BL/6 mice. The plasmid combinations injected were as follows: (1) (i) shp53 and NRAS (henceforth referred to as wt-ACVR1), (ii) shp53, NRAS, and ACVR1m. Mice were injected according to a previously described protocol (2). Briefly, plasmids were mixed in mass ratios of 1:2:2:2:2 (20 µg plasmid in a total of 40 µL plasmid mixture) with in vivo-jetPEI® (Polyplus Transfection, 201-50G) (2.8 µL per 40 µL plasmid mixture) and dextrose (5% total) and maintained at room temperature for at least 20 minutes prior to injection. Anesthesia was performed by placing the pup on ice for 2 minutes and then on a neonatal stereotaxic stage cooled to 2-8°C to maintain anesthesia. The fourth ventricle (3 mm posterior to the λ-suture and 3 mm deep) was targeted (4). Plasmid uptake and tumor development and progression was monitored as previously described (2,3). Animals were monitored daily for signs of morbidity (ataxia, impaired mobility, hunched posture, seizures, or scruffy fur). Symptomatic mice were transcardially perfused using Tyrode's solution and fixed with 4% paraformaldehyde (PFA) (2,3,5).

**Immunohistochemistry (IHC) of paraffin embedded brains**

Immunhistochemistry staining was performed as previously described (2,3). Briefly, after perfusion, mouse brains were harvested and post-fixed in 4% PFA then paraffin embedded. Tissue was sectioned using a rotary microtome. Heat mediated antigen retrieval was performed using either 1X Rodent Decloaker (Biocare Medical, RD 913) or citrate buffer (10 mM citric acid, 0.05% non-ionic detergent, pH 6) at 125°C for 30 seconds and at 90°C for 10s followed by quenching of endogenous peroxides using 0.3 % H2O2 for 30 minutes, and permeabilization with 0.025% Triton in TBS. Sections were blocked with 10% horse serum, 0.1% BSA in TBS) for 30 minutes and then incubated overnight at 4°C with primary antibody diluted in 0.1% BSA in TBS. For 3,3′-Diaminobenzidine (DAB) staining the VECTASTAIN ABC HRP Kit (Vector Laboratories, PK-4000) was used. The Alexa Flour 488 Tyramide SuperBoost Kit (Thermo Fisher Scientific,

NC1136352) was used for the phospho Smad1/Smad5/Smad8 (Ser463/465) (MilliporeSigma, AB3848-I) antibody. Bright-field images were obtained using Olympus MA BX53 microscope. Flourescent images were obtained using confocal microscopy (Carl Zeiss: MIC-System). A list of all antibodies used is included in Supplementary Table 1.

**Primary Neurospheres (NS)**

Mouse neurospheres (NS) were generated from tumors that were developed using the SB system by injection of the following plasmid combinations (1) (i) shp53 and NRAS, or (ii) shp53, NRAS, and ACVR1m into the lateral ventricle (1.5 mm AP, 0.7 mm lateral, and 1.5 mm deep from the λ-suture) following previously described protocols (2,3,5,6). When animals became symptomatic they were anesthetized then transcardially perfused with Tyrode's solution. The brains were harvested and tumors were identified by fluorescence expression. The tumor was then extracted and placed in 300 μL of media in an Eppendorf tube. Then, the tumor was mechanically dissociated using a sterile plastic pestle that fit the walls of the Eppendorf tube. This was followed by incubation with 1 mL of a cell dissociation buffer (Accutase, 423201), then filtered through a 70 μm strainer and maintained in neural stem-cell media [DMEM/F12 with L-Glutamine (Gibco, 11320-033), B-27 supplement (Gibco, 12587- 010), N-2 supplement (Gibco, 17502-048), Penicillin-Streptomycin (Corning, Cellgro, 30-001-CI), and Normocin (InvivoGen, ant-nr-1)] at 37 °C, 5% CO2. hFGF and hEGF (Shenendoah Biotech,100-26, 100-146) were supplemented twice weekly at 1 μL (20 ng/μL each stock, 1000x stock) per 1 mL media. Primary mutant ACVR1 glioma NS were transduced with pLVX-OVA to generate mACVR1-OVA NS and selected with 5 ug/mL puromycin (Sigma, P8833). pLVX-OVA was generated by cloning cytoplasmic ovalbumin from pCI-neo-cOVA (Addgene plasmid #25097) into pLVX-mCherry-c1 (Takarabio, 632561) by Nde1 and EcoR1 directed cloning.

**Western blot**

NS were harvested and re-suspended in RIPA buffer (MilliporeSigma, R0278) with 1X of Halt™ Protease and Phosphatase Inhibitor Cocktail, EDTA-free (100X) (Thermo Scientific, 78441). The Pierce™ BCA Protein Assay Kit (Thermo Scientific, 23227) was used to measure protein concentration. 20 μg of protein extract were run on a 4-12% SDS-PAGE PAGE gel (Thermo Fisher Scientific, NuPAGE®, NP0322BOX) and transferred to nitrocellulose membranes (Bio-Rad, 1620112). The membrane was probed with 1:500 of anti-phospho Smad1/Smad5/Smad8 (Ser463/465) (MilliporeSigma, AB3848-I); 1:500 of anti-Smad1 antibody (Abcam, ab63356); 1:500 of anti-Id1 (Biocheck, BCH-1/37-2); and 1:10,000 of β-actin antibody (MilliporeSigma, A1978). The secondary antibodies used were: 1:4000 of goat anti-rabbit (Dako, Agilent Technologies, P0448) and 1:20,000 of rabbit anti-mouse (Dako, Agilent Technologies, P0260 ). SuperSignal West Femto (Thermo Fisher Scientific, 34095) was used to for detection.

**In vitro experiments with ACVR1 inhibitor**

LDN-214117 is a specific ACVR1 inhibitor (7). To determine an effective concentration to inhibit ACVR1 signaling in our mouse NS we plated wild type or mutant ACVR1 cells in a T-25 flak containing growth media supplemented with varying concentrations of LDN-214117 (0.03 uM, 0.1 uM, 0.3 uM, 1 uM); Selleck Chemicals, S7147) or equivalent DMSO control. NS were incubated with inhibitor for 90 minutes and then cells were collected to assess phosphorylation of Smad1/5 and Id2 by western blot.

**Immunocytochemistry**

SU-DIPG-VI and SU-DIPG-XX1 were obtained from Dr. Michelle Monje at Stanford University (Stanford, Ca) in accordance with an institutionally approved protocol at each institution. Cells were cultured in DMEM with 10% FBS. 1 X $10^5$ cells per well were plated on glass slides coated with 2% gelatin. Cells were fixed with 4% PFA and permeabilized with PBS 1X plus 0.3% Tween. ICC was performed with 1:100 Id1 (Biocheck, BCH-1/195-14), pERK1/2 (Cell Signaling, 4370), pMEK1/2 (Santa Cruz, sc-7995-R), and 1:1000 goat anti-rabbit antibody, Alexa Flour 488 (Invitrogen, A -11034).

**RNA-seq analysis**

RNA-seq analysis was performed in collaboration with the Bioinformatics Core at the University of Michigan. Read files from the University of Michigan Sequencing Core's storage were downloaded and concatenated into a single fastq file for each sample. The quality of the raw reads data for each sample was assessed using FastQC [1] (version v0.11.3) to identify features of the data that may indicate quality problems (e.g. low quality scores, over-represented sequences, inappropriate GC content). The Tuxedo Suite software package was used for alignment, differential expression analysis, and post-analysis diagnostics. Briefly, we aligned reads to the reference genome including both mRNAs and lncRNAs (UCSC mm10) using TopHat (version 2.0.13) and Bowtie2 (version 2.2.1.). The default parameter settings for alignment were used, with the exception of: "--b2-very-sensitive" telling the software to spend extra time searching for valid alignments. We used FastQC for a second round of quality control (post-alignment), to ensure that only highquality data would be input to expression quantitation and differential expression analysis. We performed two different analysis techniques to specify differential expression: Tuxedo and DESeq2, using UCSC mm10.fa as the reference genome sequence. The volcano plot was produced using an R base script and encompasses all genes identified by our RNA-seq analysis. In order to identify significantly differentially expressed genes, we set a log2 fold change cut-off of 0.585 and –log10(FDR) greater than 1.3, where circles represent individual genes and colors as indicated (red-upregulated, green-downregulated) shown in Figure 3A. The data was analyzed using Advaita Bio's iPathwayGuide (https://www.advaitabio.com/ipathwayguide). This software analysis tool implements the 'Impact Analysis' approach that takes into consideration the direction and type of all signals on a pathway, the position, role and type of every gene, etc., as described in (8-11). Gene Ontology (GO) enrichment analysis was performed using iPathwayGuide (http://www.advaitabio.com/ipathwayguide), displaying up- and down-regulated genes associated with the same GO and was shown in Figure 3C. GO Biological Processes, selected for relevance to phenotype, were plotted in a horizontal bar graph using graphpad shown in Figure 3B. All 100 differentially expressed genes between wt-ACVR1 and mutant-ACVR1 NS were converted to excel with their corresponding gene names in uppercase in first column and log2-fold change values in second column to create the rank file. This rank file is used as an input for gene set enrichment analysis (GSEA) pre-ranking on program downloaded from Broad Institute (http://software.broadinstitute.org/gsea/index.jsp). In addition, Broad Institute provides a gene set containing all GOs (c5.all.v7.0.symbols.gmt), which is also required to perform the pre-ranking using 2000 permutations and gene set size range of 0 to 200. The enrichment map is generated using the Cytoscape platform and requires the rank and gmt files along with the positive/negative GSEA report files. Each node represents an individual GO, the size is reflective of the number of genes within that set, and the red color signifies positively regulated pathways. We set a pvalue cut off of 0.001 and FDR of 0.5. The stringency of our FDR was lowered due to encompass the underlying biological pathways differentially regulated by ACVR1 are shown in yellow in Figure 3D. The enrichment score plots for Regulation of BMP signaling Pathway and Regulation of Cell Differentiation are shown in Figure 3E and 3F respectively, which are images provided in the GSEA Pre-ranking report.

**Intratumoral injection of adenoviral vectors and radiation treatment**

We used first generation Ad.hCMV.hsFLT3L (Ad-Flt3L) + Ad.hCMV.TK (Ad-TK)(12,13). Five days post tumor implantation (dpi) mice were assigned to four treatment groups: (i) control group: empty Ad(Ad0)/saline); Ad0 was delivered intratumorally (i.t) and saline intraperitoneally (i.p.), (ii) gene therapy: Ad-TK(1×10^8 plaque-forming units (pfu)/Ad-Flt3L(2×10^8 pfu of Ad-TK) /GCV; Ad-TK/Ad-Flt3Ldelivered i.t., ganciclovir (GCV) (Biotang; RG001-1g) was administered i.p. at 25 mg/kg/daily for 10d, starting 1d post-gene therapy, (iii) Standard of care (IR): Ad0/saline+IR; an overall dose of 20 Gy IR was administered (2 Gy/d for 10d), and (iv) Gene therapy + standard of care: Ad-TK/Ad-Flt3L/GCV+IR. Intratumoral injections of adenoviral vectors was delivered in μL volume in three locations to depths of 4.4, 4.5, and 4.6 mm, at the coordinates detailed above. Sample size was n = 5 for each treatment group. For the functional analysis of T cells in the tumor microenvironment (TME), 8 days post tumor implantation mice were assigned to either the control group or the gene therapy group detailed above. Sample size was n = 3; where per sample it was necessary to pool tumor tissue from three mice for the control and five mice for the gene therapy group due to the size of the tumor.

**Flow cytometry**

For flow cytometry experiments were performed using protocols described before (12-14). Flow data were acquired on a FACSAria flow cytometer (BD Biosciences) and analyzed using Flow Jo version 10 (Treestar). Prior to staining cells with antibodies, live/dead staining was carried out using fixable viability dye (eBioscience). Then, cells were resuspended in PBS containing 2% fetal bovine serum (FBS) (flow buffer) and non-specific antibody binding was blocked with CD16/CD32 (Biolegend, 101335). All stains were carried out for 30 minutes at 4°C with 3X flow buffer washes between live/dead staining, blocking, surface staining, cell fixation, intracellular staining and data acquisition. The fixation/ permeabilization staining kit (BD Biosciences, 554714) was used for all intracellular stains. Antibody information is included in supplementary table 1. For T cell functional analysis within the TME, the cells from the tumor mass were stained with anti-mouse CD45, CD3, CD8, and SIINFEKL-H2Kb-tetramer-PE. For IFNγ stains, single cell suspensions generated from the tumor mass were stimulated with 25 ug/mL of mACVR1-OVA lysate for 24 hours in 10% FCS-containing media followed by 6 h incubation with Brefeldin and monensin. Cells were stained with CD3 and CD8 antibodies, followed by intracellular staining for IFNγ. To assess tumor cell stemness tumor cell suspensions generated from wt-ACVR1 and mACVR1 implanted tumors were stained for CD133, CD44, and Aldh1.

**Assessment of damage associated molecule release**

To assess for release of damage associated molecules after treatment with either radiation (3G) or Ad-TK (500 MOI)/GCV (25μM), 10,000 mACVR1 NS were plated on 6-well plates. The next day they were treated with Ad-TK (500 MOI) and/or radiation (3G). Following a 48 hour incubation they were treated with GCV (25 μM ). The following day they were stained with calreticulin or high mobility group box 1 (HMGB1) antibodies following flow cytometry protocols described above. To assess ATP release, we used the ATP determination kit following manufacturer's instructions (Invitrogen, A22066) to measure ATP levels in media after 48 hours of treatment. Media was used to generate the ATP standard curve.

**T cell Proliferation analysis**

Splenocytes from 21 dpi mACVR1-OVA brainstem glioma-bearing mice treated with saline or gene therapy, as detailed above, were labeled with Carboxyfluorescein succinimidyl ester (CFSE) per manufacturer's instructions and cultured with 100 nM SIINFEKL peptide (Anaspec, 60193-1; dissolved in H20 and stored in -80 °C) for 4 days. As a positive control we used splenocytes from Rag2 knockout/transgenic OT-I T cell receptor mice (Taconic, 2334) stimulated with SIINFEKL.

As a negative control we used unstimulated splenocytes. Cells were then stained with CD3 and CD8, and T cell proliferation was assessed based on CFSE dye dilution.

**Cytotoxic T cell assay**

Splenocytes from saline or gene therapy treated mice were incubated with mACVR1 NS for 24 hours at the indicated ratios (1:1, 10:1, 20:1). Lysis of tumor cells was assessed by using Annexin-V and DAPI. mACVR1 NS undergoing late apoptosis were identified as Annexin-$V^+$/$DAPI^+$ cells(15).

**Complete blood count (CBC) and serum chemistry**

Collected blood was transferred to EDTA tubes for CBC (RAM Scientific, 077058) or in serum separation tubes (Sarstedt, 41.1378.005). CBC and serum chemistry was performed by the In-Vivo Animal Core (IVAC) at the University of Michigan.

**Neuropathological analysis**

To assess the safety of vector delivery into the brainstem, non-tumor bearing animals were treated with Ad-TK/Ad-Flt3L gene therapy or saline. GCV was administered intraperitoneally one day later for 7 days. One day after the last dose of GCV was administered animals were euthanized and brains were harvested and processed for histology as detailed above. Hematoxylin and eosin (H&E) staining was performed to assess gross histopathological features (16). Immunohistochemistry staining for glial fibrillary acidic protein (GFAP) to mark astrocytes, myelin basic protein (MBP) to mark myelin sheaths and oligodendrocytes, and Iba1 to mark microglial cells.

**Hematoxylin and eosin (H&E)**

Liver tissue sections (5 µm thick) from tumor bearing animals treated with saline, radiation, gene therapy, and gene therapy and radiation were stained with H&E (16).
